# Phenotypic screening in Parkinson’s iPSC-dopamine neurons identifies the FASN-SREBP1 axis as a potential therapeutic target

**DOI:** 10.64898/2026.10.27.761977

**Authors:** Maria Claudia Caiazza, Janine Brandes, Elizabeth R. Barrow, Ricardo Márquez-Gómez, Dean Brown, Benjamin Jenkins, Uroosa Chugtai, William McGuinness, Stewart Humble, Parnaz Sharifi, Johanna L. Hoffmann, Matthieu Trigano, Albert Koulman, Ian Gurrell, Andrew Davis, David Smith, Brent Ryan, Richard Wade-Martins

**Affiliations:** Oxford Parkinson’s Disease Centre and Department of Physiology, Anatomy and Genetics, University of Oxford; South Park Road, Oxford, OX1 3QU, UK; Kavli Institute for Nanoscience Discovery, University of Oxford; Dorothy Crowfoot Hodgkin Building, South Park Road, Oxford, OX1 3QU, UK; Open Innovation, Hit Discovery, Discovery Sciences, BioPharmaceuticals R&D, AstraZeneca, Boston, US; Wellcome Trust-MRC Institute of Metabolic Science Metabolic Research Laboratories, Cambridge, CB2 0QQ, UK; Open Innovation, Hit Discovery, Discovery Sciences, BioPharmaceuticals R&D, AstraZeneca, Cambridge, UK

## Abstract

Parkinson’s disease (PD) remains highly challenging to model in vitro, particularly when seeking robust functional phenotypes for target discovery. Patient-derived induced pluripotent stem cell dopaminergic neurons (iPSC-DANs) offer a promising platform to address this challenge. Because calcium dyshomeostasis is well-described in PD pathophysiology, it offers a platform for phenotypic screening. We have previously shown that *GBA-N370S* iPSC-DANs display impaired intracellular calcium release. Here, we used this disease-relevant phenotype to undertake a 4,108-compound phenotypic screen in *GBA-N370S* (*GBA-NS*) iPSC-DANs. This screen identified fatty acid synthase (FASN) as the most enriched protein target. The ability of compounds to rescue calcium release correlated with biochemical potency against FASN across multiple chemical series. We confirmed that FASN knockdown increased intracellular calcium release, supporting an on-target effect. Untargeted lipidomics demonstrated broad remodelling of the neuronal lipidome after FASN inhibition, confirming target engagement. FASN inhibition also increased active β-glucocerebrosidase (GCase), reduced extracellular α-synuclein release and ER stress, and altered RASGRF1 expression. Mechanistically, FASN inhibition activated sterol regulatory element-binding protein 1 (SREBP1), and SREBP1 overexpression alone was sufficient to increase intracellular calcium release, identifying a FASN-SREBP1-calcium homeostasis axis in human neurons. Finally, the FASN inhibitor BI 99179 achieved brain exposure and reduced brain fatty acid levels while inducing an SRE-associated transcriptional response in a Parkinson’s mouse model, providing evidence of in vivo target engagement. These findings establish patient iPSC-neuron functional screening as a tractable discovery platform to identify targets which can be taken in vivo and nominate the FASN-SREBP1 axis as a candidate therapeutic pathway in PD.

**One Sentence Summary:** We applied a disease-relevant phenotypic screen in patient neurons to uncover a lipid-calcium regulatory axis that rescues multiple pathogenic pathways in Parkinson’s.

## INTRODUCTION

Parkinson’s disease (PD) is the second most common neurodegenerative disorder and is characterised by α-synuclein aggregation and the preferential vulnerability of A9 dopaminergic neurons (DANs) in the substantia nigra pars compacta (SNpc). Despite its growing clinical burden, PD remains incurable, highlighting the need for experimental systems that capture early pathogenic mechanisms and support therapeutic discovery. Patient-derived induced pluripotent stem cell (iPSC) models provide a platform to study disease-relevant cellular phenotypes directly in vulnerable human neurons. Extensive phenotypic, transcriptomic, proteomic and metabolomic analyses have demonstrated that iPSC models recapitulate key aspects of PD biology, underscoring their relevance (*1, 2*). Beyond disease modelling, iPSC-based systems offer a powerful opportunity for phenotypic drug screening directly in human patient neurons, an approach that remains underexplored in neurodegenerative disease (*3–5*).

Genetic studies have strongly implicated the lysosomal-autophagic pathway in PD pathogenesis (*6, 7*). Among PD-associated genes, *GBA1*, which encodes the lysosomal enzyme GCase, represents the most common strong genetic risk factor. Heterozygous *GBA1* mutations, including N370S and L444P, reduce GCase activity and disrupt lysosomal homeostasis (*8*). Notably, decreased GCase activity has also been observed in synucleinopathy models lacking *GBA1* mutations, supporting a broader role for GCase dysfunction in PD (*9*). Consistent with this, iPSC-DANs carrying *GBA1* mutations exhibit a range of disease-relevant phenotypes, including lyswosomal perturbation, mitochondrial dysfunction, synaptic abnormalities, and defects in intracellular calcium handling. Both our previous work and that of others have identified impaired organelle calcium handling in *GBA*-PD iPSC-DANs, highlighting calcium dyshomeostasis as a robust and functionally relevant phenotype in this system (*10, 11*).

Calcium dyshomeostasis is well described as a key feature of PD pathophysiology (*12–14*). We previously demonstrated that *GBA-NS* iPSC-DANs exhibit reduced ionomycin-induced intracellular calcium release, indicating lower organellar calcium (*11*). Here, we exploited this phenotype to establish a phenotypic screening platform in patient-derived neurons and screened a well-annotated library of 4,108 compounds covering more than 1,800 human targets.

The screen identified FASN as the most enriched target class associated with rescue of impaired intracellular calcium release. FASN is a key enzyme in de novo fatty acid synthesis and has recently emerged as a PD-relevant gene, and its inhibition has been reported to ameliorate PINK1 deficiency-associated phenotypes in PD models (*15–17*). Using complementary pharmacological and genetic approaches, we validated FASN as a modulator of intracellular calcium release in human iPSC-DANs. Consistent with target engagement, untargeted lipidomic profiling revealed broad remodelling of the neuronal lipidome after FASN inhibition. FASN inhibition also improved multiple PD-relevant phenotypes, including GCase activity, ER stress and extracellular α-synuclein release. Mechanistically, FASN inhibition activates SREBP1, and SREBP1 overexpression is sufficient to enhance intracellular calcium release in iPSC-DANs. Finally, *in vivo* studies with the compound BI 99179 provide initial evidence that pharmacological FASN inhibition can engage the pathway in the brain. Overall, our work identifies a FASN-SREBP1-calcium cellular axis as a candidate therapeutic pathway in PD. Together, these findings establish patient iPSC-based phenotypic screening as a tractable discovery platform with the potential to identify therapeutic targets which can be validated in vivo to de-risk the drug discovery process.

## RESULTS

### Phenotypic screening in PD patient iPSC-DANs

We performed a high-throughput small-molecule screen to correct the ionomycin-induced intracellular calcium release phenotype in *GBA-NS* iPSC-DANs (*11*). To enable efficient phenotypic screening, we scaled our iPSC differentiation protocol up to T175 flasks to generate sufficient midbrain floor plate progenitors (mFPPs) for screening and miniaturised the calcium assay from 96- to 384-well format (Figure 1A). The disease-associated reduction in ionomycin-induced intracellular calcium release was retained in both 96- and 384-well formats (Figure 1B and 1C), and the *GBA-NS* iPSC-DANs used in screening showed robust phenotype in the miniaturised assay (Figure 1D). Neuronal identity was maintained following miniaturisation, with comparable TH-positive cell proportions between plate formats (Figure S1A and S1B).

**Fig. 1.**
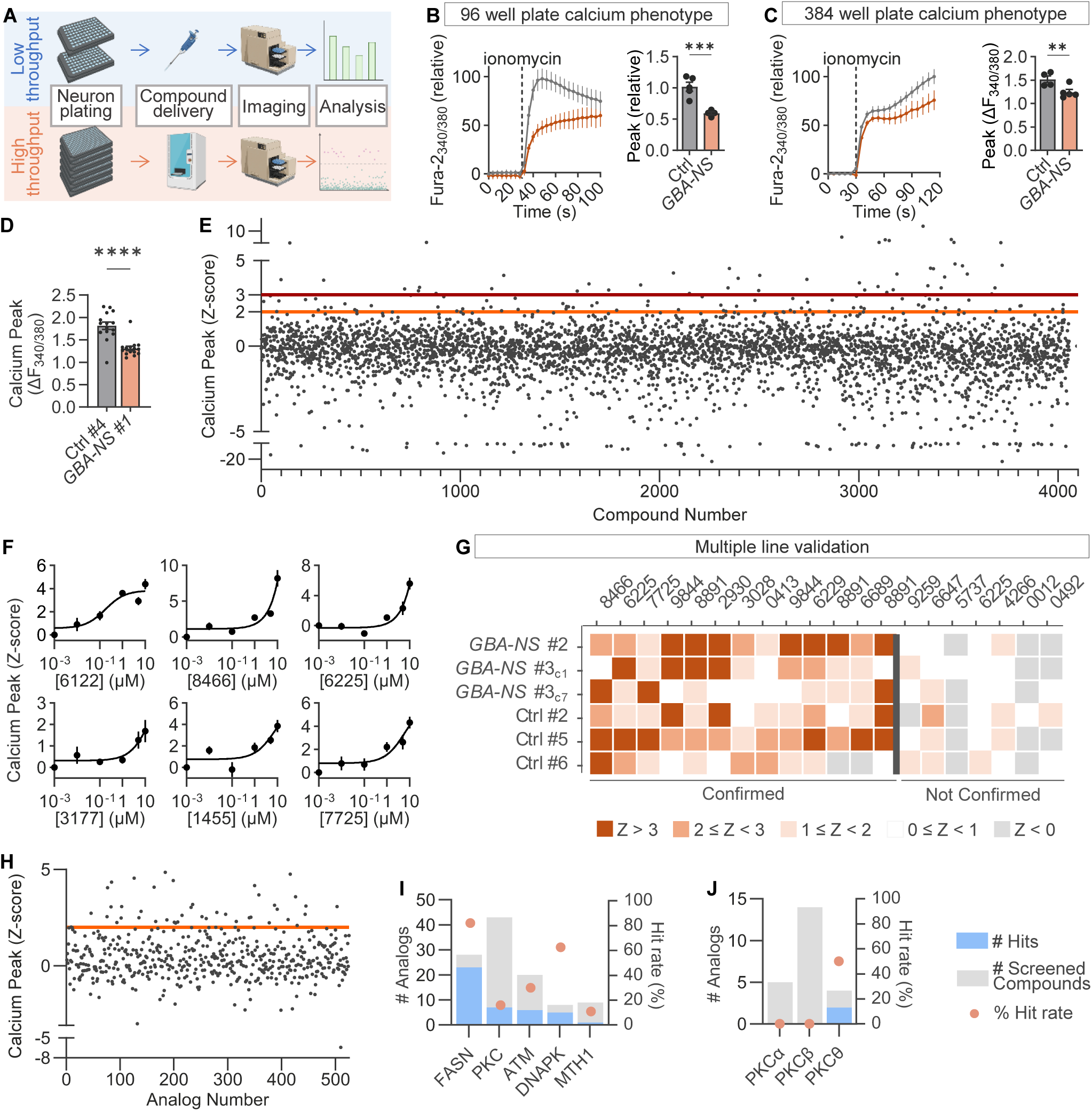
Phenotypic screening in patient iPSC-DANs identifies modulators of ionomycin-induced organelle calcium release. (A) Schematic of the low-throughput and high-throughput screening workflows, comprising neuron plating, compound delivery, calcium imaging and analysis. (B) Ionomycin-evoked Fura-2 calcium responses and peak-amplitude quantification in Ctrl and *GBA-NS* iPSC-DANs in 96-well format. Normalised Fura-2 AM ratiometric traces were averaged across lines and differentiations; peak amplitudes are shown relative to Ctrl. Statistical analysis by one-tailed unpaired t-test. (C) Ionomycin-evoked calcium-response quantification in Ctrl and *GBA-NS* iPSC-DANs in 384-well format. Normalised Fura-2 AM ratiometric traces were averaged across lines taken from one representative differentiation. Statistical analysis by one-tailed unpaired t-test. (D) Peak calcium-response amplitude in the Ctrl #4 and *GBA-NS* #1 lines selected for screening in 384-well format. (E) Primary phenotypic screen of 4,108 compounds in *GBA-NS* #1 iPSC-DANs. Calcium peak amplitudes are shown as Z-scores; orange and red lines indicate Z = 2 and Z = 3 thresholds, respectively. (F) Representative concentration-response validation of six primary screening hits. (G) Multi-line validation of selected hits across *GBA-NS* and Ctrl iPSC-DAN lines. Colours indicate calcium peak-amplitude Z-score; compounds to the left of the divider were confirmed across the validation criteria. (H) Calcium peak-amplitude Z-scores for 525 analogues of confirmed primary-screen targets; the orange line indicates Z = 2. (I) Number of hits, number of analogues screened and percentage hit rate for analogues targeting FASN, PKC, ATM, DNAPK and MTH1. (J) Deconvolution of PKC inhibitors by isoform, showing the number screened, number of hits and hit rate for PKCα, PKCβ and PKCθ.

For the screen, we selected 4,108 compounds from an AstraZeneca phenotypic screening library, each with sub-100 nM potency against its primary target, spanning more than 1,800 human targets. Screening was performed in iPSC-DANs derived from a patient carrying the *GBA-NS* mutation (referred to as *GBA-NS* #1) and showing a representative deficit in ionomycin-induced intracellular calcium release. Differentiated neurons were treated with compounds at 10 µM for 48 h in triplicate wells and assessed for phenotypic rescue.

We identified 123 compounds (a 3% hit rate) that increased ionomycin-induced intracellular calcium release in *GBA-NS* #1 (Figure 1E). These results demonstrate that disease-relevant phenotypic screening can be achieved at scale in patient-derived iPSC-DANs, despite the variability inherent to neuronal screening assays (Figure S1C and S1D).

Primary screening hits were validated by concentration-dependent modulation of intracellular calcium release in *GBA-NS* #1 (Figure 1F). To assess generalisability, compounds were tested across iPSC-DANs from multiple donors, including *GBA-NS* PD patients and healthy controls, confirming effects across multiple genetic backgrounds (Figure 1G). Overall, we confirmed 23 compounds as hits (Z ≥ 2) acting in a concentration-dependent manner and in multiple lines of iPSC-DANs.

Confirmed hit compounds were mapped to their annotated protein targets, identifying 17 distinct proteins among the 23 compounds (Table S1). Based on this analysis, we screened 525 analogues targeting these proteins, of which 44 (8.4%) increased calcium release (Z ≥ 2) (Figure 1H). Target-level analysis identified several enriched classes, including FASN, PKC, ATM, DNAPK and MTH1, with FASN showing the highest hit rate (Figure 1I). Further interrogation of PKC inhibitors implicated PKCθ as the predominant PKC isoform associated with calcium modulation (Figure 1J).

### FASN inhibition increases ionomycin-induced intracellular calcium release

Given compound analogue screening identified FASN as the most enriched target, 110 FASN analogues were screened, of which 78 and 38 increased calcium release with Z-scores >1 and >2, respectively (Figure 2A). The majority of these compounds (n=96) clustered into five chemically distinct classes, with multiple clusters demonstrating increased calcium release (Figure 2B and 2C). To further validate the link between FASN inhibition and calcium release, we correlated the strength of the calcium release modulation of 38 analogues with their biochemical potency against FASN. This revealed a strong association between FASN inhibition (pIC50) and intracellular calcium release (r = 0.52, p < 0.0001, Spearman’s correlation) (Figure 2D).

**Fig. 2.**
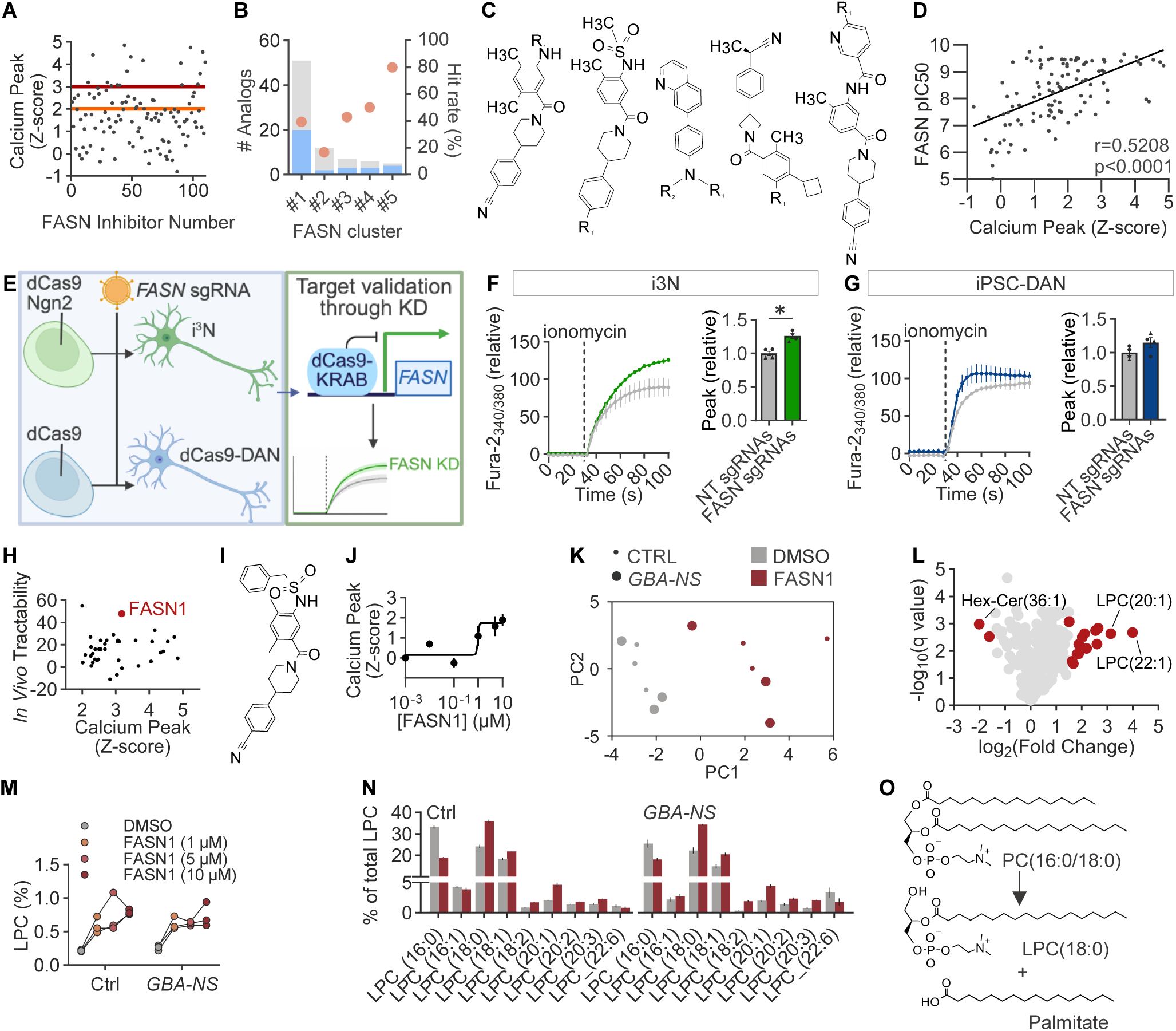
Identification, genetic validation and target-engagement characterisation of FASN inhibition. (A) Calcium peak-amplitude Z-scores across 110 FASN inhibitors tested in *GBA-NS* iPSC-DANs; orange and red lines indicate Z = 2 and Z = 3, respectively. (B) Distribution of FASN analogues across five chemical clusters, showing the number of analogues tested and percentage hit rate for each cluster. (C) Representative chemical structures from the five FASN inhibitor clusters. (D) Relationship between biochemical FASN potency (pIC50) and ionomycin-induced calcium peak-amplitude Z-score (r = 0.5208, P < 0.0001; Spearman correlation). (E) Schematic of CRISPRi-mediated FASN target validation. (F) Ionomycin-evoked calcium responses and peak-amplitude quantification in i3N transduced with non-targeting or FASN sgRNAs. Normalised Fura-2 AM ratiometric traces were averaged across two sgRNAs; peak amplitudes are shown relative to non-targeting controls. (G) Ionomycin-evoked calcium responses and peak-amplitude quantification in dCas9 iPSC-DANs transduced with non-targeting or FASN sgRNAs. (H) Prioritisation of FASN inhibitors by cellular calcium activity and *in vivo* tractability, highlighting FASN1. (I) Chemical structure of FASN1. (J) Concentration-dependent effect of FASN1 on ionomycin-induced calcium release in *GBA-NS* iPSC-DANs. (K) Principal component analysis of lipidomic profiles from Ctrl and *GBA-NS* iPSC-DANs treated with DMSO or FASN1 (5 µM). (L) Volcano plot of lipid changes following FASN1 treatment, highlighting significantly altered lipid species including Hex-Cer(36:1), LPC(20:1) and LPC(22:1). (M) Overall LPC levels in Ctrl and *GBA-NS* iPSC-DANs following treatment with DMSO or FASN1 (1, 5 or 10 µM). (N) Distribution of individual LPC species as a percentage of total LPC in Ctrl and *GBA-NS* iPSC-DANs treated with DMSO or FASN1 (5 µM); data are mean ± SEM. (O) Schematic representation of phosphatidylcholine hydrolysis generating LPC and free palmitate. *P ≤ 0.05.

To validate FASN as the target responsible for modulating calcium release, we used CRISPR interference (CRISPRi) (Figure 2E). We first used the well-established glutamatergic GT3-dCas9 i3 neuron (i3N) system (*18, 19*). Two independent FASN sgRNAs reduced FASN RNA and protein abundance (Figure S2A-C) and increased ionomycin-induced intracellular calcium release compared with non-targeting controls (Figure 2F). Transduction efficiency is shown in Figure S2D.

We next generated a stable polyclonal CRISPRi iPSC line using a piggyBac transposon system expressing nuclear-localised dCas9-KRAB and an mNeonGreen reporter (Figure S2E). The engineered line retained dopaminergic differentiation capacity and reporter expression (Figure S2F, S2G and S2I), and sgRNA transduction efficiency was quantified in differentiated dCas9-DANs (Figure S2H). FASN knockdown was confirmed by western blot (Figure S2J and S2K) and increased ionomycin-induced calcium release in iPSC-DANs (Figure 2G), recapitulating the effect observed in i3N. Together, these genetic data support FASN as the on-target mediator of the calcium phenotype.

### Lipidomic profiling confirms FASN target engagement by FASN1

To prioritise a FASN inhibitor for further mechanistic studies, compounds were ranked by comparing an in vivo tractability score, integrating chemical properties, inter-species FASN inhibition potency, pharmacokinetics and MDCK transport, with cellular activity in the calcium assay (Figure 2H). FASN1 was selected as the lead compound (Figure 2I) and was shown to increase ionomycin-induced intracellular calcium release in *GBA-NS* iPSC-DANs in a concentration-dependent manner (Figure 2K), with minimal toxicity across the tested concentration range (Figure S2L).

FASN catalyses the de novo synthesis of palmitate, a key precursor for complex lipids, and its inhibition is expected to reshape the neuronal lipidome. We therefore performed untargeted lipidomic profiling of iPSC-DANs treated with FASN1 or DMSO (Figure 2H). Principal component analysis (PCA) revealed clear separation between FASN1-treated and vehicle-treated samples (Figure 2K), with treatment-dependent separation also evident across the concentration series (Figure S2M).

Differential lipid analysis showed broad remodelling following FASN1 treatment, with several lysophosphatidylcholine (LPC) species among the most strongly altered lipids (Figure 2L). Profiling of LPC composition further demonstrated an increase in the total LPC and substantial redistribution of individual LPC species in both Ctrl and *GBA-NS* iPSC-DANs (Figure 2M and 2N). Notably, while the total LPC was increased, the proportional abundance of palmitate-containing LPC(16:0) was reduced following FASN inhibition, consistent with mobilisation of palmitate-containing phosphatidylcholine (PC) to generate LPC and free palmitate (Figure 2O). This remodelling is consistent with a homeostatic response to altered palmitate availability and supports on-target FASN inhibition, in line with previous reports (*20*).

### FASN inhibition improves PD-relevant phenotypes and calcium homeostasis

Consistent with previous reports (*10, 21–23*), active GCase levels were reduced in *GBA-NS* iPSC-DANs compared with Ctrl neurons (Figure 3A). The specificity and dynamic range of the MDW941 activity-based GCase probe were validated using the GCase inhibitor CBE (Figure S3A-C).

**Fig. 3.**
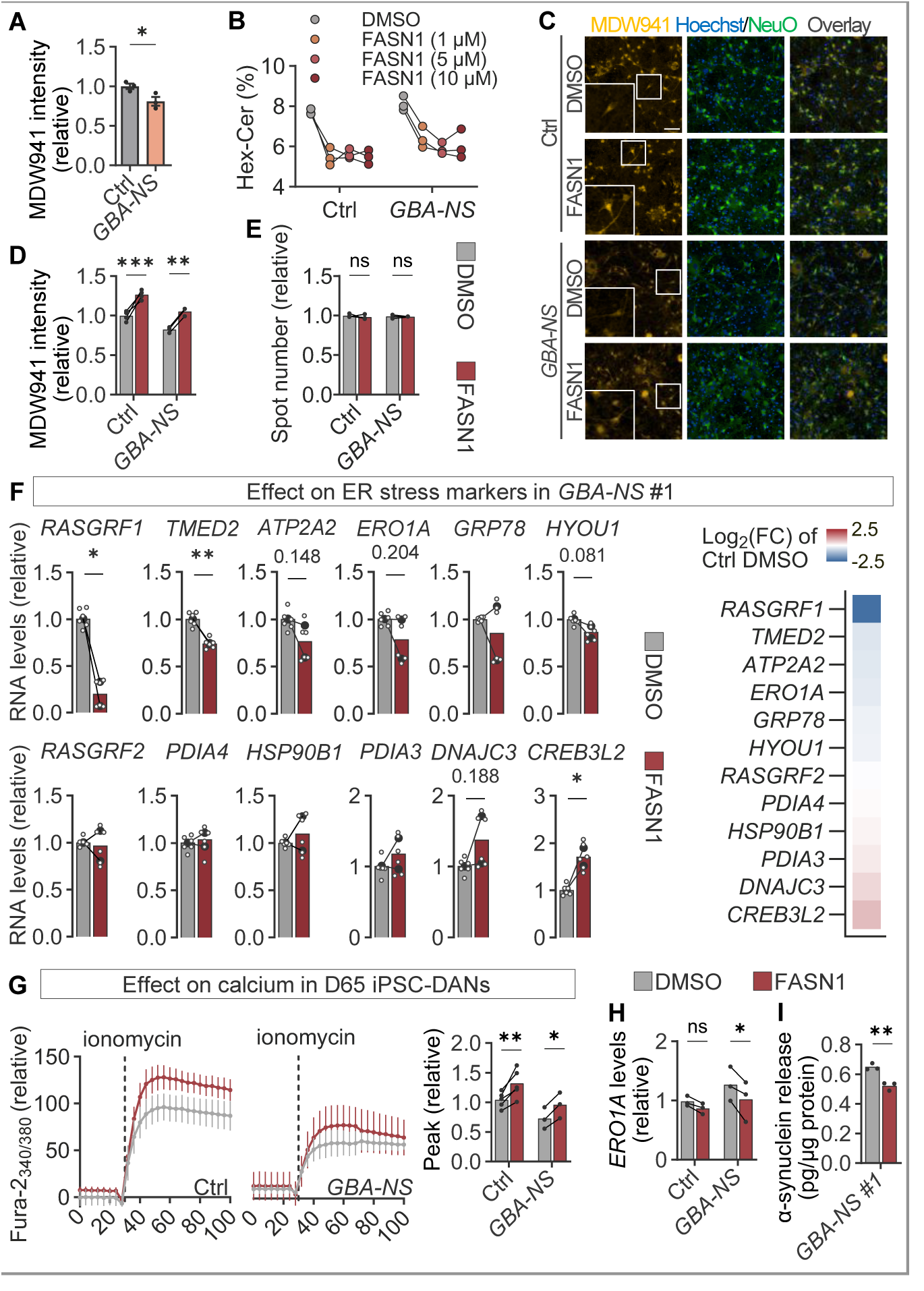
FASN inhibition improves PD-relevant phenotypes and calcium homeostasis in *GBA-NS* iPSC-DANs. (A) Active GCase signal is reduced in *GBA-NS* relative to Ctrl iPSC-DANs, measured using the fluorescent activity-based probe MDW941. Each dot represents an individual line mean; data are mean ± SEM. Statistical analysis by one-tailed Mann-Whitney test. (B) Hex-Cer levels in Ctrl and *GBA-NS* iPSC-DANs following treatment with DMSO or FASN1 (1, 5 or 10 µM), determined by lipidomics. (C) Representative images of Ctrl and *GBA-NS* neurons treated with DMSO or FASN1 and stained with Hoechst (blue), NeuO (green) and MDW941 (yellow). Scale bar, 50 µm. (D) Quantification of MDW941 corrected spot intensity following FASN1 (5 μM) treatment. (E) Quantification of MDW941-positive spot number. For D-E, each dot represents an individual line mean across independent differentiations; data are mean ± SEM. Statistical analysis by paired two-way ANOVA with Fisher’s LSD multiple-comparisons test. (F) mRNA expression of UPR effectors and chaperones in *GBA-NS* #1 iPSC-DANs following DMSO or FASN1 (10 μM) treatment. Each solid dot represents a differentiation averaged across three technical replicates (hollow dots); lines link data belonging to the same differentiation; bars are means. Statistical analysis by Nested t-test. (G) Normalised Fura-2 AM ratiometric traces showing ionomycin-evoked calcium release in mature DIV65 Ctrl and *GBA-NS* iPSC-DANs treated with DMSO or FASN1 (10 μM), with quantification of peak amplitudes, expressed relative to Ctrl DMSO. Statistical analysis by two-way ANOVA. (H) ERO1A mRNA expression at D65 in Ctrl and *GBA-NS* iPSC-DANs treated with DMSO or FASN1 (10 μM). (I) Extracellular α-synuclein release from *GBA-NS* #1 DANs following DMSO or FASN1 (10 μM) treatment, measured by MSD assay. Each dot represents a technical replicate; data are mean ± SEM. Statistical analysis by two-tailed Student’s t-test. Statistical analysis by two-way ANOVA. *P < 0.05; **P < 0.01; ***P < 0.001.

In the lipidomic dataset, FASN inhibition with FASN1 decreased levels of the GCase substrate hexosylceramide (Hex-Cer) in a concentration-dependent manner (Figure 3B). In the lipidomic dataset, levels of the GCase substrate hexosylceramide (Hex-Cer) decreased in a concentration-dependent manner following FASN inhibition with FASN1 (Figure 3B). Consistent with this reduction, FASN1 significantly increased the intensity of the active GCase signal in both Ctrl and *GBA-NS* iPSC-DANs (Figures 3C and 3D), without altering the number of MDW941-positive puncta (Figure 3E). These findings suggest FASN1 enhanced GCase trafficking and activity, rather than increased GCase synthesis or lysosome biogenesis. In line with this interpretation, qPCR analysis of genes within the CLEAR pathway revealed no significant changes following FASN1 treatment (Figure S3D).

Because ER stress is an established feature of *GBA-NS* iPSC-DANs (*21, 24, 25*) and could potentially be alleviated by improved GCase trafficking (*26*), we next investigated whether FASN inhibition modulated this phenotype. FASN1 selectively remodelled the adaptive unfolded protein response (UPR) in *GBA-NS* #1 iPSC-DANs (Figure 3F). Notably, RASGRF1, a calcium-responsive regulator of Ras signalling, was markedly reduced following FASN1 treatment. This occurred concomitantly with the restoration of intracellular calcium levels, suggesting that altered *RASGRF1* expression may accompany the normalisation of calcium-dependent signalling. *ATP2A2*, which encodes the ER calcium pump SERCA2, also showed a trend towards decreased expression, potentially reflecting an adaptive response to restored calcium homeostasis. In addition, *ERO1A* showed a trend towards decreased expression. Of particular interest, *ERO1A* was previously identified as one of the prominently dysregulated ER stress-associated genes in *GBA-NS* iPSC-DANs (*25*). As ER stress and UPR dysregulation emerge as later phenotypes in *GBA-NS*iPSC-DANs (*25*), we therefore extended neuronal maturation from day 35 to day 65 (D65) to determine whether the effect of FASN1 on *ERO1A* became more pronounced under conditions of greater cellular stress. We first confirmed that FASN1 retained its ability to rescue the calcium phenotype at D65 (Figure 3G) and subsequently assessed *ERO1A* expression. At this later time point, FASN1 significantly reduced *ERO1A* expression in *GBA-NS* iPSC-DANs (Figure 3H).

Finally, we sought to investigate the effect of FASN inhibition on α-synuclein pathology in *GBA-NS,* and observed that FASN reduced extracellular α-synuclein release in the *GBA-NS* #1 screening line (Figure 3I).

### FASN inhibition activates SREBP1 and demonstrates target engagement in vitro and in vivo

Fatty acids are known to inhibit the lipogenic transcription factor SREBP1 through multiple mechanisms (*27–30*). Consistent with this, SREBP1 protein abundance increased in both Ctrl and *GBA-NS* iPSC-DANs following FASN1 treatment (Figure 4A and 4B). To determine whether this increase was accompanied by an SREBP-associated transcriptional response, we measured a panel of established SRE-regulated genes and PD-relevant candidate genes. FASN1 altered the expression of multiple genes in both Ctrl and *GBA-NS*iPSC-DANs, consistent with activation of an SRE-associated transcriptional programme (Figure 4C). FASN protein abundance was also increased after FASN1 treatment, consistent with feedback regulation of the pathway (Figure 4D and 4E).

**Fig. 4.**
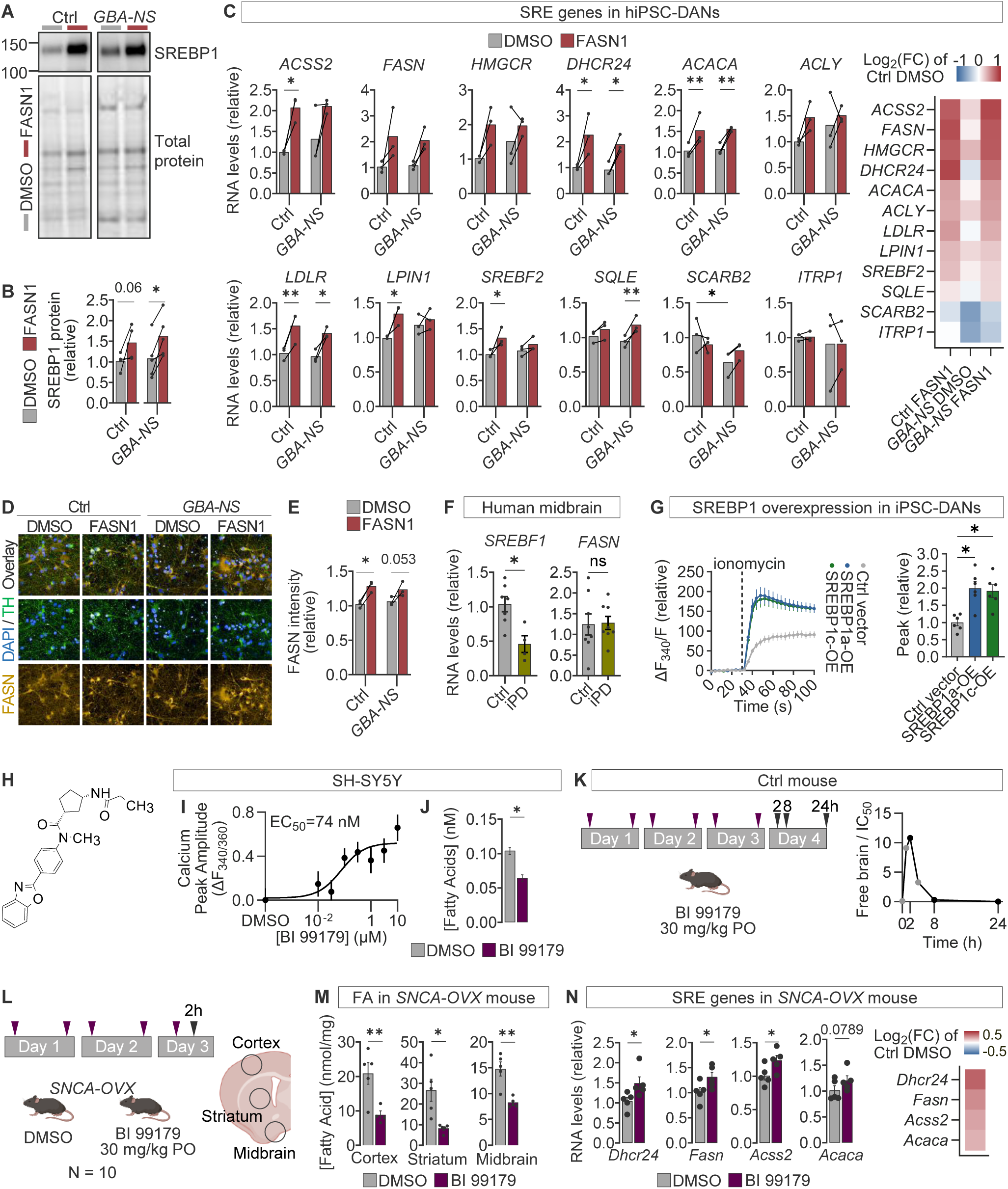
FASN inhibition activates SREBP1 signalling and demonstrates *in vivo* target engagement. (A-B) Western blot and quantification showing SREBP1 protein abundance following FASN1 (5 µM) treatment in Ctrl and *GBA-NS* iPSC-DANs. SREBP1 signal was normalised to total protein; each dot represents a line mean from independent differentiations, and data are mean ± SEM. Statistical analysis by one-tailed Wilcoxon test. (C) Expression of SRE-associated genes (*ACLY*, *ACSS2*, *DHCR24*, *FASN*, *HMGCR*, *ITPR1, LDLR*, *LPIN1*, *PLCG2, SCARB2, SREBF2*, and *SQLE*) in Ctrl and *GBA-NS* iPSC-DANs treated with DMSO or FASN1 (5 µM). Bar plots show relative RNA levels, and the heatmap shows log_2_ fold change relative to Ctrl DMSO. (D) Representative images of FASN (yellow), TH (green) and DAPI (blue) in Ctrl and *GBA-NS* iPSC-DANs treated with DMSO or FASN 1 (5 µM). (E) Quantification of FASN fluorescence intensity from D. Statistical analysis by two-way ANOVA (F) SREBF1 and FASN mRNA levels in human midbrain tissue from healthy Ctrl and idiopathic PD donors. Each dot represents an individual donor mean from technical replicates; data are mean ± SEM. Statistical analysis by one-tailed Wilcoxon test. (G) Ionomycin-evoked calcium responses and peak-amplitude quantification in iPSC-DANs overexpressing SREBP1a or SREBP1c compared with control vector. Traces are shown as normalised ΔF340/F; each dot represents a line mean. Statistical analysis by Friedman test followed by BKY multiple-comparison procedure (Q = 0.05). (H) Chemical structure of BI 99179. (I) Concentration-response for BI 99179 on ionomycin-induced calcium peak amplitude in SH-SY5Y cells (EC50 = 74 nM). (J) Measurement of FA levels in response to BI 99179 treatment in SH-SY5Y cells. Statistical analysis by one-tailed Welch’s t-test. (K) BI 99179 dosing and tissue-collection paradigm and free-brain BI 99179 exposure relative to mouse FASN IC50 over 24 h in wild-type mice. Black points represent experimental data and grey points represent predicted data (L) BI 99179 dosing and tissue-collection paradigm in *SNCA*-OVX mice; BI 99179 was administered at 30 mg/kg PO and cortex, striatum and midbrain were collected 2 h after the final administration. (M) Fatty acid levels in cortex, striatum and midbrain from DMSO- and BI 99179-treated *SNCA*-OVX mice. Statistical analysis by one-tailed Welch’s t-test for each brain region. (N) Expression of SRE-associated genes in SNCA-OVX mouse brain following DMSO or BI 99179 treatment. Bar plots show relative RNA levels, and the heatmap shows log_2_ fold change relative to DMSO. Statistical analysis by one-tailed lognormal Welch’s t-test for each gene. *P < 0.05; **P < 0.01; ns, not significant.

Previous studies have suggested that SREBP1 and FASN may be directly implicated in PD (*16, 31*). We therefore measured SREBF1 and FASN mRNA in human midbrain tissue from healthy controls and individuals with idiopathic PD. SREBF1 expression was reduced in PD samples, whereas FASN expression was unchanged (Figure 4F). To test whether SREBP1 activation is sufficient to recapitulate the calcium effect of FASN inhibition, we overexpressed the two SREBP1 isoforms (SREBP1a and SREBP1c) in iPSC-DANs using lentiviral constructs (*32*). Overexpression of either isoform (Figure S4A-C) increased ionomycin-induced intracellular calcium release compared with a control vector (Figure 4G), demonstrating that increased SREBP1 activity is sufficient to phenocopy the calcium effect of FASN inhibition.

To explore the translational tractability of FASN inhibition, we next used the compound BI 99179 (*33*) (Figure 4H). In SH-SY5Y cells, BI 99179 increased ionomycin-induced calcium release in a concentration-dependent manner (EC50 = 74 nM; Figure 4I) and reduced cellular fatty acid levels (Figure 4J).

Pharmacokinetic analysis in wild-type mice demonstrated measurable free brain exposure relative to FASN IC50 (mouse IC50 = 80 nM) over 24 h in mice following the dosing and collection paradigm described (Figure 4K). Black points represent experimental data, and grey points represent predicted brain data based on measured free plasma concentration (Figure S4D) and BI 99179 brain Kp,uu of 0.13. We therefore treated the Parkinson’s *SNCA*-OVX transgenic mouse model with BI 99179 (30 mg/kg, PO) or vehicle according to the dosing and collection paradigm shown in Figure 4L. The treatment paradigm did not affect the weight of the treated mice (Figure S4E). BI 99179 reduced fatty acid levels in cortex, striatum and midbrain (Figure 4M). In parallel, BI 99179 increased expression of several SRE-associated genes, including *Acss2*, *Dhcr24*, *Fasn* and *Acaca* (Figure 4N), consistent with the in vitro data in the iPSC-DAN model of FASN inhibition. Together, these data provide evidence of central target engagement and activation of a compensatory SRE-associated transcriptional response following FASN inhibition in vivo.

## DISCUSSION

Here, we have demonstrated that phenotypic screening in human iPSC-DANs is a tractable strategy for therapeutic discovery in PD, and that the FASN-SREBP1-calcium signalling axis represents a candidate therapeutic pathway in the disease. Using *GBA-NS* iPSC-DANs, we show that pharmacological inhibition and genetic knockdown of FASN consistently increase ionomycin-induced calcium release, supporting a direct role for FASN in regulating this disease-relevant phenotype. Lipidomic profiling confirmed target engagement and revealed broad remodelling of the neuronal lipidome following FASN inhibition. FASN inhibition also improved multiple PD-relevant phenotypes, including GCase activity, extracellular α-synuclein release and ER stress. Mechanistically, our data support a model in which FASN inhibition activates SREBP1, and increased SREBP1 activity is sufficient to enhance intracellular calcium release. The link between SREBP1 pathway upregulation and calcium handling is likely to be multifactorial, potentially involving changes in ER or lysosomal calcium handling, including IP3R-dependent calcium release, lipid-dependent remodelling of organelle membranes, including altered membrane contact-site biology (*34–36*).

Finally, studies with BI 99179 extend these observations in vivo, demonstrating brain exposure, reduced brain fatty acid levels and an SRE-associated transcriptional response in the Parkinson’s *SNCA*-OVX mouse model. Together, these findings position the FASN-SREBP1 axis as a point of convergence between lipid metabolism, calcium signalling, lysosomal function and cellular stress responses, and support FASN/SREBP as potential therapeutic targets.

An emerging body of evidence supports a broader role for lipid dysregulation in neurodegeneration. PD has recently been described as a “fatty acidopathy”, reflecting widespread alterations in fatty acid and lipid homeostasis that may influence α-synuclein aggregation and cellular vulnerability (*15*). Human genetic data further support the relevance of this pathway, with recent large-scale multi-ancestry GWAS identifying the *FASN* locus as a susceptibility region for PD, and other studies identifying *SREBF1* as a risk locus in sporadic PD (*16, 31*). In this context, our findings align with prior work showing that FASN inhibition rescues PINK1-associated phenotypes across multiple model systems, including patient-derived neurons, in part through remodelling mitochondrial lipids such as cardiolipin and improving mitochondrial function (*17*). Similarly, SREBP1 has been shown to be important for mitophagy induction (*37*). Our findings extend this growing body of evidence by implicating the FASN-SREBP1 axis in calcium homeostasis and lysosomal function. Together, these observations suggest that modulation of fatty acid synthesis may have pleiotropic effects on key PD-relevant pathogenic pathways.

The relevance of FASN inhibition may extend beyond PD. In Alzheimer’s disease models, increased FASN expression and elevated palmitate levels have been linked to lipid peroxidation and neurodegeneration, while pharmacological inhibition of FASN has been shown to remodel lipid metabolism, reduce inflammation, and improve cognitive outcomes (*38*). These findings, together with our data, support the concept that targeting lipid metabolic pathways may represent a broader therapeutic strategy across neurodegenerative disorders.

Certain limitations to this study exist. First, although our data support a role for SREBP1 activation in mediating the effects of FASN inhibition on intracellular calcium release, we did not dissect the molecular pathways linking lipid remodelling to improved calcium homeostasis. Defining these pathways will be important, particularly to identify narrower-acting targets downstream of FASN-SREBP1. Second, it will be important to determine the broader effects of FASN inhibition on neuronal function, including its impact on other PD-relevant pathways and on mechanisms implicated in other neurodegenerative diseases.

Several steps will be needed before these findings can be translated clinically. The BI 99179 mouse data provide initial evidence that pharmacological FASN inhibition in the brain can generate measurable target engagement; however, the dose-response relationship, durability of pathway engagement, consequences of chronic FASN inhibition and therapeutic efficacy on disease-relevant endpoints remain to be established. Given the availability of FASN inhibitor chemical matter from oncology and metabolic disease programs, there may be opportunities to accelerate translational development. Stratification of patients by genetic background, including *GBA1*, *PINK1*, *FASN* and *SREBP1* status, may also help identify groups most likely to benefit from this approach.

Overall, our findings identify FASN as a regulator of calcium homeostasis and lysosomal function in human iPSC-DANs and support the FASN-SREBP1 axis as a candidate therapeutic pathway in PD. More broadly, our work demonstrates the potential of human iPSC-based phenotypic screening to uncover disease-relevant biology and accelerate target discovery in neurodegeneration.

## MATERIALS and METHODS

### Key resources table

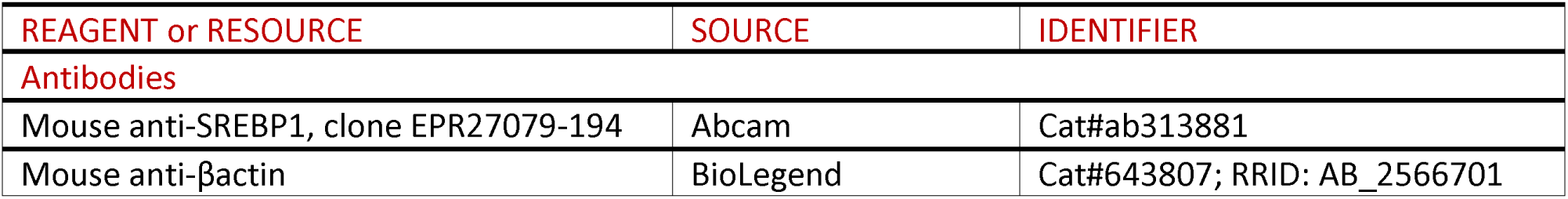
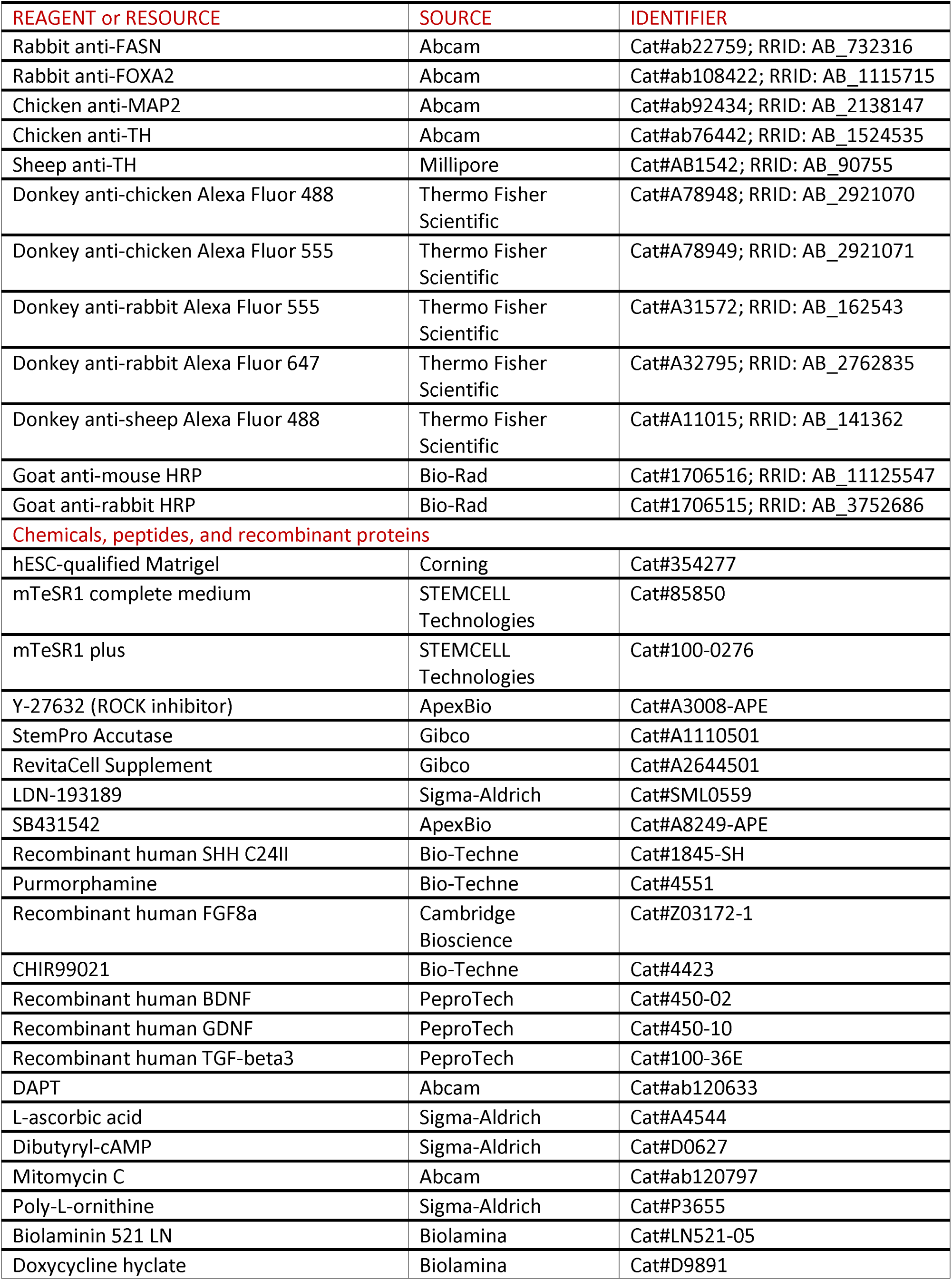
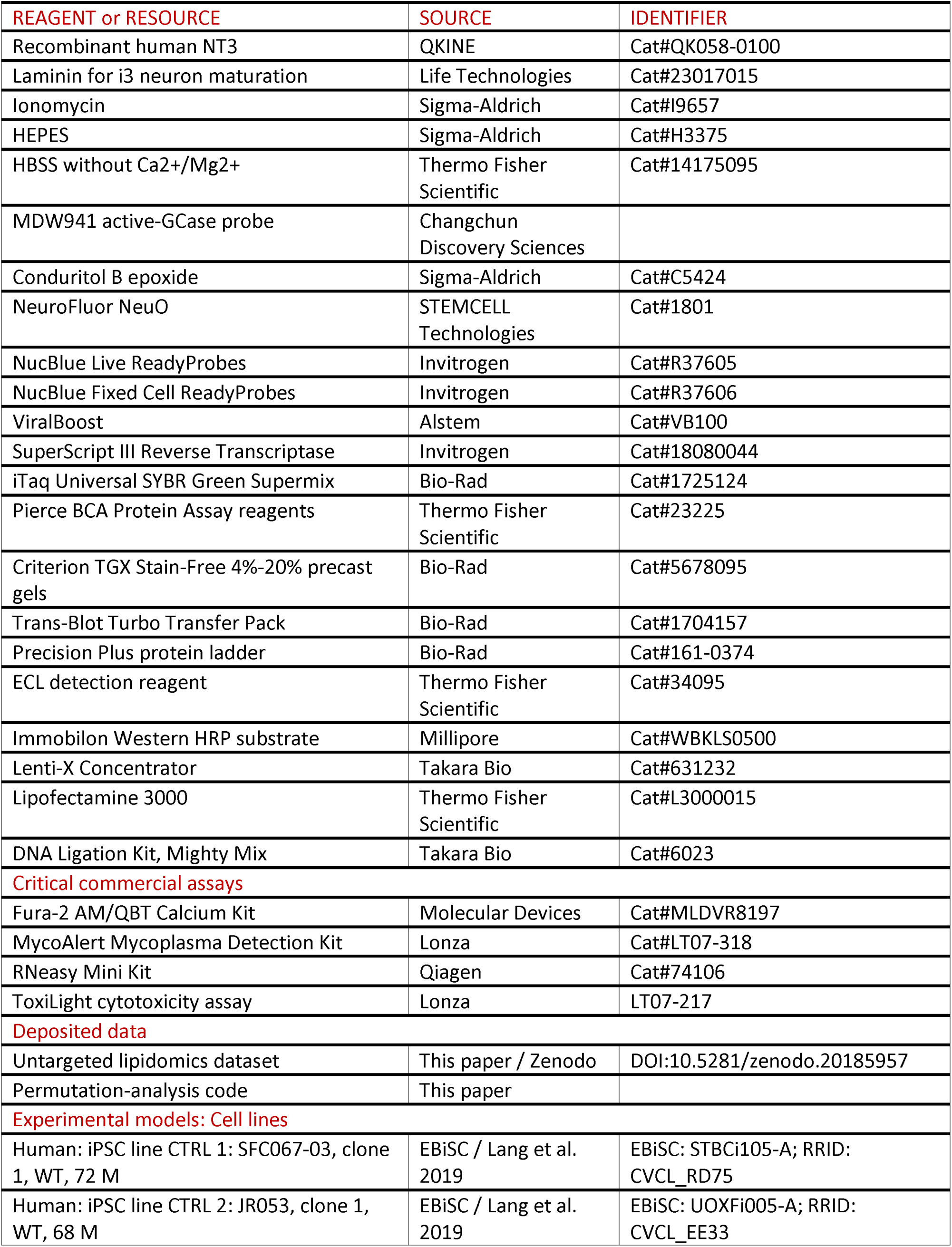
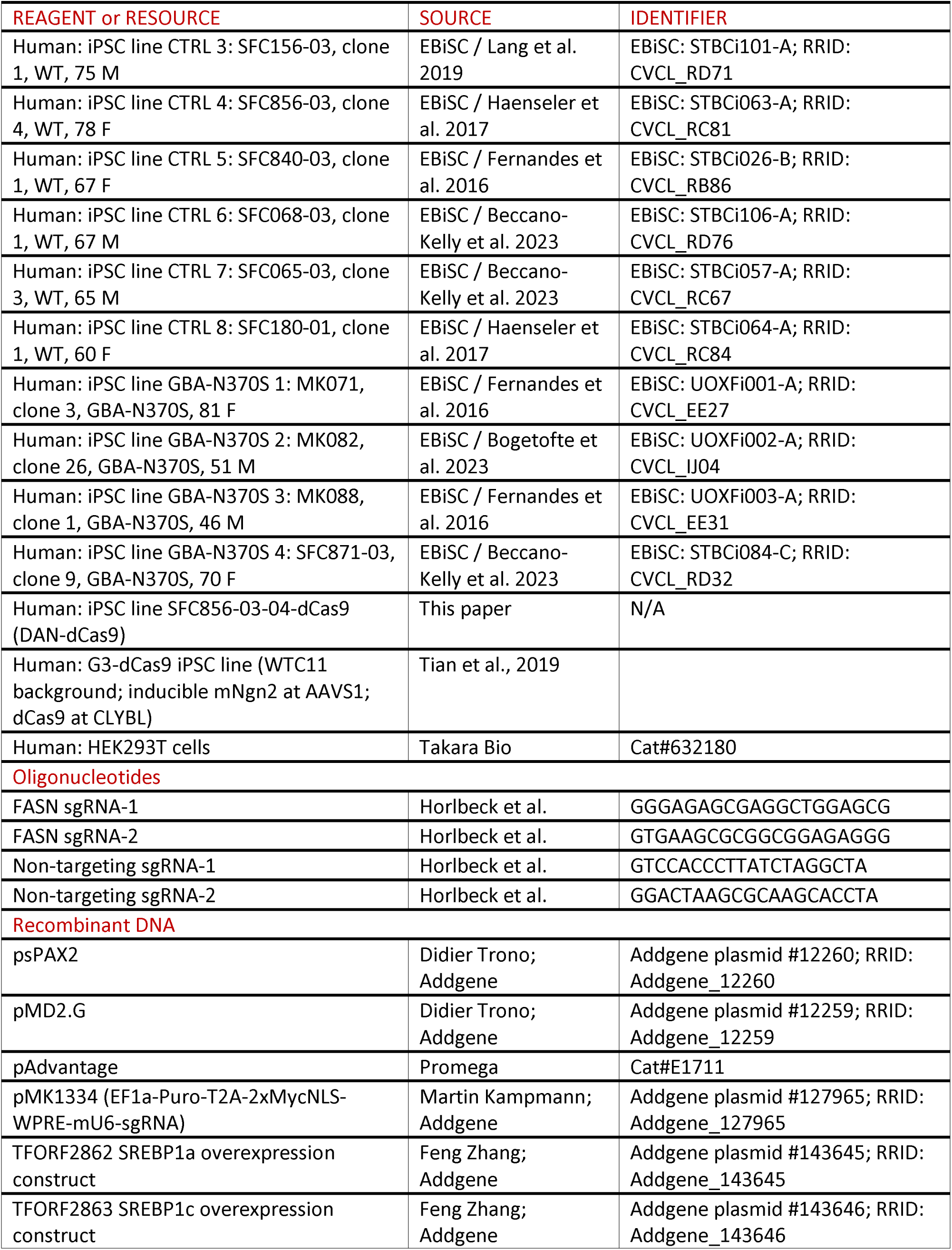
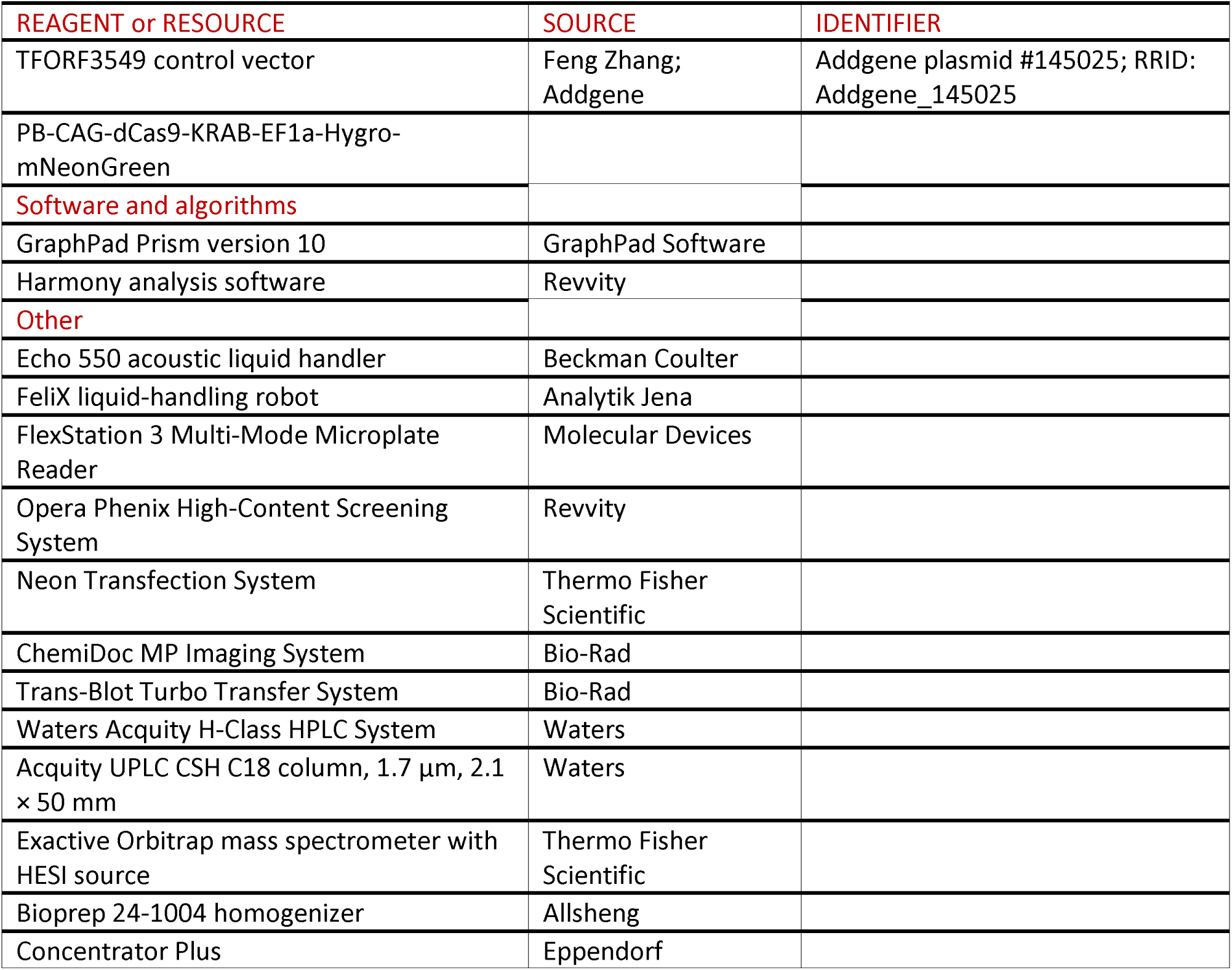

## METHOD DETAILS

### iPSC maintenance and quality control

iPSCs were maintained on hESC-qualified Matrigel-coated plates (Corning, Cat#354277) in mTeSR1 medium (STEMCELL Technologies, Cat#85850) at 37°C and 5% CO₂. Colonies were passaged at approximately 70% confluence using 0.5 mM EDTA. Cultures were routinely tested negative for mycoplasma using the MycoAlert assay (Lonza, Cat#LT07-318) and monitored for chromosomal abnormalities by pseudokaryotyping.

### Generation of the DAN-dCas9 CRISPRi iPSC line

iPSCs were pre-treated with Y-27632 (ApexBio, Cat#A3008-APE; 10 µM, 1 h), dissociated with Accutase, and resuspended at 3 × 10^7^ cells/mL. A total of 3 × 10^5^ cells was mixed with 200 ng PiggyBac transposase plasmid and 400 ng PB-CAG-dCas9-KRAB-EF1α-Hygro-mNeonGreen in Neon Buffer R. Cells were nucleofected in 10 µL tips using the Neon Transfection System (1,400 V, 20 ms, one pulse), plated onto Matrigel-coated 24-well plates in mTeSR1 containing 1% RevitaCell, and allowed to recover.

Hygromycin selection was initiated at 70%-80% confluence in four 48 h bursts separated by recovery periods (125 µg/mL for bursts 1-2 and 250 µg/mL for bursts 3-4). Polyclonal selected cells were expanded, cryopreserved, and differentiated without clonal isolation. The PiggyBac transposase plasmid and PB-CAG-dCas9-KRAB-EF1α-Hygro-mNeonGreen plasmid were gifts from Michael Ward; sequence maps are available from the lead contact upon request. Following selection, approximately 90% of NucBlue-counterstained nuclei were mNeonGreen-positive.

### Differentiation of iPSCs into dopaminergic neurons

Dopaminergic neurons were generated as previously described (*39*) with introduction of the expansion of midbrain floor-plate progenitors (mFPPs) (*40, 41*). The day-by-day formulation used in the thesis is consolidated below.

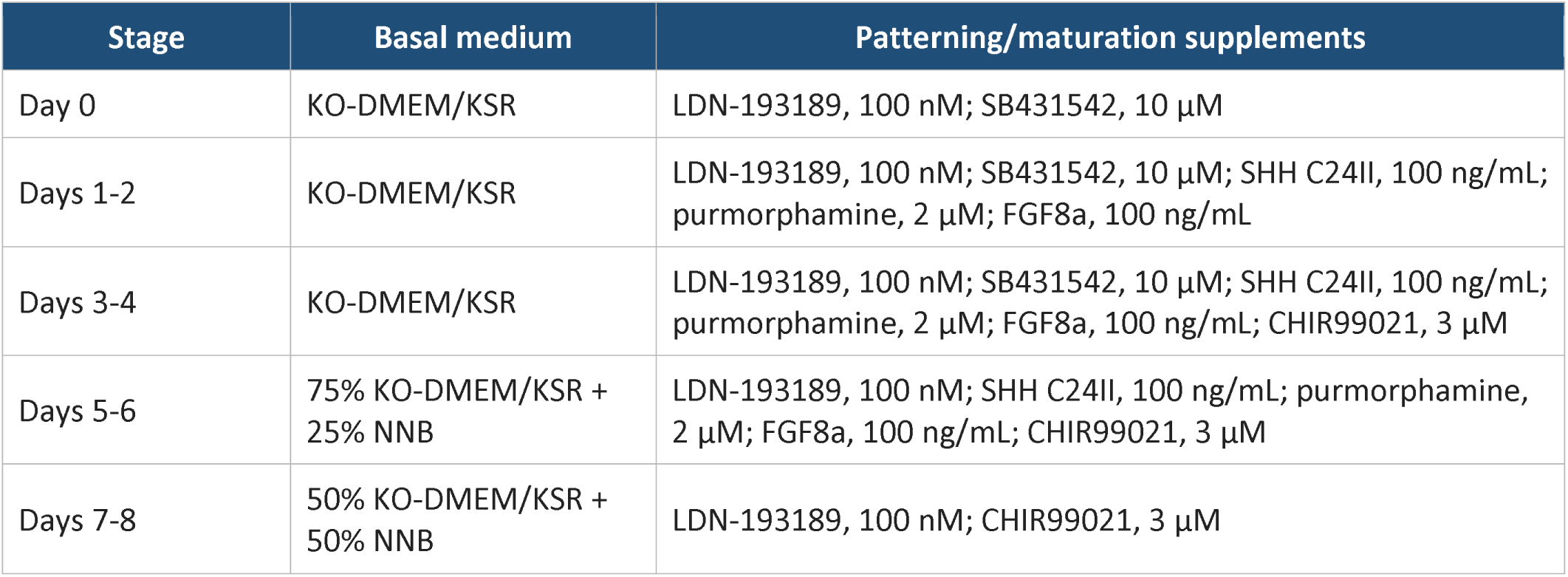
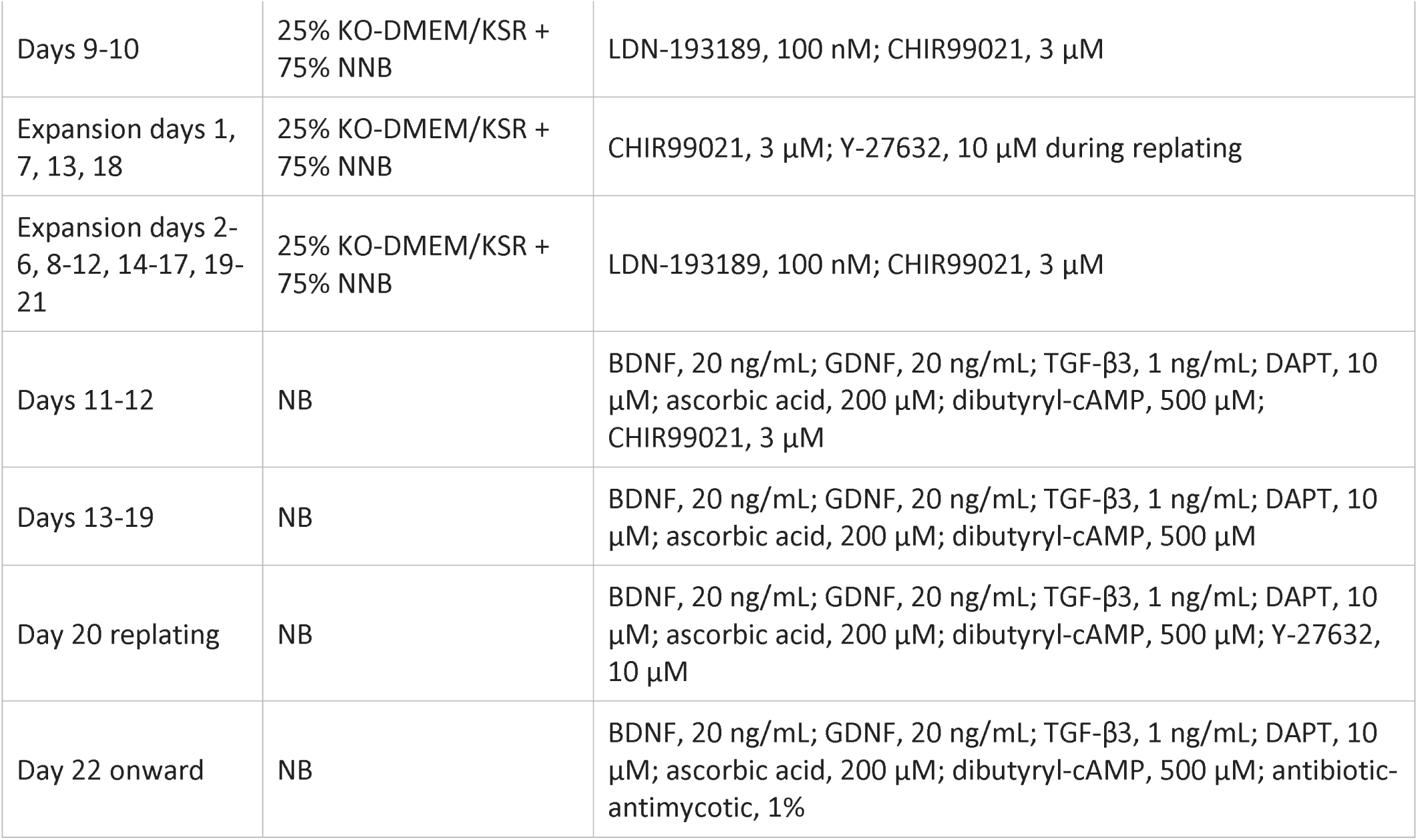

| Stage | Basal medium | Patterning/maturation supplements |
|---|---|---|
| Day 0 | KO-DMEM/KSR | LDN-193189, 100 nM; SB431542, 10 µM |
| Days 1-2 | KO-DMEM/KSR | LDN-193189, 100 nM; SB431542, 10 µM; SHH C24II, 100 ng/mL; purmorphamine, 2 µM; FGF8a, 100 ng/mL |
| Days 3-4 | KO-DMEM/KSR | LDN-193189, 100 nM; SB431542, 10 µM; SHH C24II, 100 ng/mL; purmorphamine, 2 µM; FGF8a, 100 ng/mL; CHIR99021, 3 µM |
| Days 5-6 | 75% KO-DMEM/KSR + 25% NNB | LDN-193189, 100 nM; SHH C24II, 100 ng/mL; purmorphamine, 2 µM; FGF8a, 100 ng/mL; CHIR99021, 3 µM |
| Days 7-8 | 50% KO-DMEM/KSR + 50% NNB | LDN-193189, 100 nM; CHIR99021, 3 µM |
| Days 9-10 | 25% KO-DMEM/KSR + 75% NNB | LDN-193189, 100 nM; CHIR99021, 3 µM |
| Expansion days 1, 7, 13, 18 | 25% KO-DMEM/KSR + 75% NNB | CHIR99021, 3 µM; Y-27632, 10 µM during replating |
| Expansion days 2-6, 8-12, 14-17, 19-21 | 25% KO-DMEM/KSR + 75% NNB | LDN-193189, 100 nM; CHIR99021, 3 µM |
| Days 11-12 | NB | BDNF, 20 ng/mL; GDNF, 20 ng/mL; TGF-β3, 1 ng/mL; DAPT, 10 µM; ascorbic acid, 200 µM; dibutyryl-cAMP, 500 µM; CHIR99021, 3 µM |
| Days 13-19 | NB | BDNF, 20 ng/mL; GDNF, 20 ng/mL; TGF-β3, 1 ng/mL; DAPT, 10 µM; ascorbic acid, 200 µM; dibutyryl-cAMP, 500 µM |
| Day 20 replating | NB | BDNF, 20 ng/mL; GDNF, 20 ng/mL; TGF-β3, 1 ng/mL; DAPT, 10 µM; ascorbic acid, 200 µM; dibutyryl-cAMP, 500 µM; Y-27632, 10 µM |
| Day 22 onward | NB | BDNF, 20 ng/mL; GDNF, 20 ng/mL; TGF-β3, 1 ng/mL; DAPT, 10 µM; ascorbic acid, 200 µM; dibutyryl-cAMP, 500 µM; antibiotic-antimycotic, 1% |

KO-DMEM/KSR consisted of 82% KnockOut DMEM, 15% KnockOut Serum Replacement, 1% 2-mercaptoethanol, 1% L-glutamine, and 1% MEM non-essential amino acids. NNB consisted of 97.5% Neurobasal medium, 0.5% N2 supplement, 1% B-27 supplement without vitamin A, and 1% L-glutamine. NB consisted of 97% Neurobasal medium, 2% B-27 supplement without vitamin A, and 1% L-glutamine. iPSCs were single-cell passaged on day -2 and plated at 1.5 × 10^6^ cells per well in 6-well plates. From day 0, cultures in 6-well plates received 4 mL medium per well. Full medium changes were performed on days 0, 1, 3, 5, 7, 9, 11, 13, 15, 17, and 19 and on alternating expansion days; 50% medium changes were performed on intervening days. mFPPs were dissociated with Accutase and passaged 1:2 on expansion days 1, 7, 13, and 18. Depending on experimental requirements, mFPPs were used without expansion or expanded for up to 21 days (3 weeks).

For final replating, assay plates were coated overnight with 0.01% poly-L-ornithine and then overnight with biolaminin (10 µg/mL). Cells were dissociated with Accutase for 5-7 min at 37°C, centrifuged at 350 × g for 5 min, counted, and plated at the densities below. On day 22, cultures were exposed to mitomycin C (1 µg/mL, 1 h), washed by complete medium replacement, and subsequently received 50% medium changes three times weekly. For CRISPRi experiments, cells were replated at DIV16 to improve transduction efficiency.

| Plate format | Cells/well | Plating volume | Primary use |
|---|---|---|---|
| 384-well | 25,000 | 15 µL | Calcium imaging |
| 384-well | 20,000 | 12 µL | High-content imaging |
| 96-well half-area | 35,000 | 21 µL | Calcium imaging |
| 96-well half-area | 27,000 | 16 µL | High-content imaging |
| 96-well | 85,000 | 50 µL | Calcium imaging |
| 96-well | 60,000 | 35 µL | High-content imaging |
| 12-well | 1.7 × 106 | 1 mL | RNA/protein extraction |
| 6-well | 4 × 106 | 2.5 mL | Protein extraction |

Assays were generally performed at approximately DIV35 and at DIV65 where indicated.

### Differentiation of G3-dCas9 iPSCs into i3 neurons

G3-dCas9 iPSCs were differentiated by doxycycline-inducible expression of Ngn2 as previously described (*19*). During induction, cells were maintained on Matrigel in DMEM/F12 with HEPES (Gibco, Cat#11330032) supplemented with N2, non-essential amino acids, and L-glutamine, each at 1:100. Doxycycline hyclate (Sigma-Aldrich, Cat#D9891) was added at 2 µg/mL for the first 3 days of induction. At DIV3, cells were replated onto 0.01% poly-L-ornithine at 4 × 10^4^ cells per well in half-area 96-well plates or 5 × 10^5^ cells per well in 24-well plates and transduced with sgRNA lentivirus. Cells were matured in medium containing BDNF (10 ng/mL), NT3 (10 ng/mL), and laminin (1 µg/mL). A full medium change was performed after replating, followed by 50% medium changes twice weekly. No puromycin selection was applied. Differentiation quality was assessed by neuronal morphology. Assays were performed at DIV10.

### Compound handling and treatment

Compounds were prepared as 10 mM stocks in DMSO and stored at room temperature under a nitrogen atmosphere in Roylan StoragePods. Compounds were arrayed by acoustic transfer using an Echo 550 and dispensed onto neuronal cultures with a FeliX liquid-handling robot. Unless otherwise stated, cells were treated for 48 h before assay, and the final DMSO concentration was 0.1% (v/v) in all wells, including vehicle controls.

### Phenotypic screen

A total of 4,108 compounds from an AstraZeneca phenotypic screening library were tested in GBA-N370S #1 iPSC-DANs. Compounds were applied at 10 µM for 48 h in triplicate wells, after which ionomycin-induced intracellular calcium release was measured by Fura-2 imaging. Hits were defined at a Z-score ≥ 2.

Primary hits were re-tested at 10⁻^3^, 10⁻^2^, 10⁻^1^, 1, and 10 µM (0.001-10 µM; a five-point, 10-fold dilution series) in *GBA-NS* #1 iPSC-DANs and at single dose (10 µM) across multiple control and *GBA-NS* donor lines. Twenty-three compounds were considered confirmed hits based on concentration-dependent activity and Z ≥ 2 in multiple lines. Confirmed compounds were mapped to 17 annotated protein targets. A follow-up set of 525 analogues directed against these targets was screened at 10 µM, followed by a focused set of 110 FASN analogues screened at 10 µM.

### HEK293T cells

HEK293T cells (Takara, Cat#632180) below passage 15 were used for lentiviral production. Cells were cultured in high-glucose DMEM with GlutaMAX (Gibco, Cat#61965059) supplemented with 10% fetal bovine serum (Sigma-Aldrich, Cat#F7524), without antibiotics, and were routinely tested for mycoplasma.

### Lentiviral production, concentration and titration

HEK293T cells below passage 15 were co-transfected with psPAX2 (Addgene #12260), pMD2.G (Addgene #12259), pAdvantage (Promega, Cat#E1711), and the relevant lentiviral transfer plasmid using Lipofectamine 3000. After 24 h, the medium was replaced with high-glucose DMEM with GlutaMAX (Gibco, Cat#61965059) containing 10% FBS (Sigma-Aldrich, Cat#F7524) and ViralBoost. Viral supernatant was harvested 48 h after the medium change, clarified by low-speed centrifugation and passage through a 0.45 µm filter, concentrated approximately 100-fold with Lenti-X Concentrator, resuspended in PBS, aliquoted, and stored at -80°C.

Fluorescently tagged viruses were titrated in 384-well plates by acoustic transfer to generate a dilution series, followed by transduction and imaging on an Opera Phenix high-content system. sgRNA viruses were dosed to achieve approximately 60% transduction efficiency. Virus was added directly to the culture medium at replating and removed by medium exchange after 1 day for i3 neurons and 2 days for iPSC-DANs. For non-tagged viruses the control titration was used as proxy.

### sgRNA cloning and FASN CRISPR interference

Annealed oligonucleotides encoding 20-nt sgRNA targeting sequences were ligated into BstXI/BlpI-digested pMK1334 (EF1a-Puro-T2A-2xMycNLS-WPRE-mU6-sgRNA; Addgene #127965) using DNA Ligation Kit, Mighty Mix. Chemically competent E. coli were transformed, ampicillin-resistant colonies were isolated, and the sgRNA cassette was confirmed by Sanger sequencing. Two independent FASN-targeting guides and two non-targeting controls were used.

### SREBP1 overexpression

SREBP1a (TFORF2862; Addgene #143645), SREBP1c (TFORF2863; Addgene #143646), and control vector (TFORF3549; Addgene #145025) lentiviral constructs were used. Lentivirus was generated as above and applied to iPSC-DANs.

### Fura-2 AM calcium imaging

Cells were loaded with Fura-2 AM using the Fura-2 QBT Calcium Kit (Molecular Devices, Cat#MLDVR8197) in HBSS without Ca2+ or Mg2+ supplemented with 20 mM HEPES (pH 7.4) for 60-90 min at 37°C, as previously described (*42*). Fluorescence was acquired on a FlexStation 3 at excitation wavelengths of 340 and 380 nm and an emission wavelength of 510 nm. For 96-well plates, each well was read every 4 s for 100 s. For 384-well plates, each well was read every 6 s for 120 s. Ionomycin was injected at 30 s to a final concentration of 5 µM.

The ratiometric signal was calculated as R = F340/F380. Baseline was defined as the mean of reads 1-8, and peak amplitude as the maximum signal through read 26 minus baseline. Peak amplitudes were expressed relative to matched control wells or lines. In order to avoid spectral overlap between the EGFP in TFORF3549, for the SREBP overexpression experiment, only the calcium bound wavelength of Fura-2 was recorded (F340) and data was normalised by ΔF/F0.

For CRISPRi experiments and SREBP1 overexpression, wells with a peak amplitude below 50% of the plate median were excluded. No non-response threshold was applied in the primary compound screen. For representation, all normalised traces were further normalised as (F-F0)/Fmax ∗ 100, where Fmax is the per-well post-ionomycin maximum, enabling cross-differentiation comparison. Peak amplitudes were then expressed relative to matched controls unless otherwise stated.

### Cell-toxicity assay

Cytotoxicity of FASN1 was evaluated by measuring adenylate kinase released into conditioned medium using the ToxiLight cytotoxicity assay (Lonza, Cat#LT07-217).

### Active GCase imaging with MDW941

MDW941 was custom-synthesized for this study by Changchun Discovery Sciences, Ltd, based on a previously published protocol (*43*). iPSC-DANs were incubated with MDW941 (5 nM) in maturation medium for 16 h at 37°C and 5% CO₂. For an inhibitor control, cells were pre-treated with conduritol B epoxide (Sigma-Aldrich, Cat#C5424; 1 mM, 1 h) before MDW941 addition. Neurons were counterstained with NeuroFluor NeuO (STEMCELL Technologies, Cat#1801), and nuclei were counterstained with NucBlue Live ReadyProbes (Thermo Fisher Scientific, Cat#R37605). Cells were gently washed with pre-warmed HBSS containing Ca2+ and Mg2+ and imaged live on an Opera Phenix using identical acquisition settings across conditions.

### Extracellular alpha-synuclein release

Total extracellular alpha-synuclein was measured using the MSD The U-PLEX Human α-Synuclein Kit according to manufacturer’s instruction. Data was normalised to total cell protein levels measured by BCA.

### RNA extraction, reverse transcription and qPCR

Total RNA was isolated using the RNeasy Mini Kit and reverse-transcribed with SuperScript III according to the manufacturers’ instructions. qPCR was performed using iTaq Universal SYBR Green Supermix with 40 cycles of 95°C for 15 s, 60°C for 25 s, and 72°C for 20 s. Melt-curve analysis was used to verify a single amplification product. Relative expression was calculated using the 2⁻ΔΔCt method.

Primer sequences are provided in the Supplemental Table 2.

### Western blotting

Cells were sonicated on ice for 20 s in RIPA buffer (50 mM Tris, pH 7.5; 150 mM NaCl; 1% NP-40; 0.5% sodium deoxycholate; 0.1% SDS) supplemented with protease inhibitor. Total protein was quantified by BCA assay. Samples were prepared in 6× Laemmli buffer and boiled at 95°C for 5 min for non-membrane-bound targets. 15-30 µg of each sample were loaded on the gel.

Proteins were separated on 4%-20% Criterion TGX Stain-Free gels, the gel was UV-activated for total-protein visualization, and proteins were transferred to PVDF with the Trans-Blot Turbo system.

Membranes were blocked in 5% milk in TBST for 1 h, incubated overnight at 4°C with rabbit anti-FASN (1:2,000; Abcam, Cat#ab22759), mouse anti-SREBP1 clone EPR27079-194 (1:1,000; Abcam, Cat#ab313881), or Direct-Blot HRP anti-β-actin (1:50,000; BioLegend, Cat#643807), washed three times for 5 min, incubated with the appropriate HRP-conjugated secondary antibody for 1 h at room temperature, washed, and detected by ECL on a ChemiDoc MP.

### Immunocytochemistry and high-content analysis

Cells were fixed in 4% paraformaldehyde in PBS for 10 min at room temperature, permeabilized with 0.1% Triton X-100 in PBS containing sodium azide for 6 min, and blocked for 1 h in 10% normal donkey serum in 0.1% Triton X-100/PBS-azide. Primary antibodies were diluted in blocking buffer and incubated overnight at 4°C. Cells were washed twice in PBS and incubated for 50-90 min at room temperature with fluorophore-conjugated secondary antibodies and NucBlue Fixed Cell ReadyProbes, protected from light. Cells were washed twice and stored in PBS-azide before imaging.

### Lipidomics

The protein-precipitation liquid extraction protocol described previously (*37*). Briefly, 650bµL of chloroform was added to each sample, followed by thorough mixing. The samples were then homogenised using a Bioprep 24-1004 homogeniser (Allsheng, Hangzhou City, China) run at speed: 4.5 m/s, time: 30 seconds for 2 cycles. Then 100 µL of the LIPID internal standard (1-10 µM in methanol), 100 µL of the CARNITINE internal standard (5bµM in methanol) and 150 µL of methanol was added to each sample, followed by thorough mixing. Then, 400bµL of acetone was added to each samples. The samples were vortexed and centrifuged for 10bminutes at ∼20,000 g to pellet any insoluble material. The supernatant was pipetted into separate 2bmL screw cap amber-glass auto-sampler vials (Agilent Technologies, Cheadle, United Kingdom). The organic extracts were dried down to dryness using a Concentrator Plus system (Eppendorf, Stevenage, United Kingdom) run for 60 minutes at 60°C.

For lipidomics LC-MS analysis: the samples were reconstituted in 100 µL of 2:b1: 1 (propan-2-ol, acetonitrile and water, respectively) then thoroughly vortexed. The reconstituted sample was transferred into a 250 μL low-volume vial insert inside a 2 mL amber glass auto-sample vial ready for liquid chromatography with mass spectrometry detection (LC-MS) lipidomics analysis. Full chromatographic separation of intact lipids was achieved using Waters Acquity H-Class HPLC System (Waters, Hertfordshire, United Kingdom) with the injection of 10 µL onto a Waters Acquity UPLC® CSH C18 column; 1.7 µm, I.D. 2.1 mm X 50 mm, maintained at 55 degrees Celsius. Mobile phase A was 6:4, acetonitrile and water with 10 mM ammonium formate. Mobile phase B was 9:1, propan-2-ol and acetonitrile with 10 mM ammonium formate. The flow was maintained at 500bµL per minute through the following gradient: 0.00bminutes_40% mobile phase B; 1.5 minutes_40% mobile phase B; 8.00 minutes_99% mobile phase B; 10.00 minutes_99% mobile phase B; 10.10 minutes_40% mobile phase B; 12.00 minutes_40% mobile phase B. The sample injection needle was washed using 9:1, propan-2-ol and acetonitrile [strong wash] and 2: 1: 1 (propan-2-ol, acetonitrile and water) [weak wash]. The mass spectrometer used was the Thermo Scientific Exactive Orbitrap with a heated electrospray ionisation source (Thermo Fisher Scientific, Hemel Hempstead, United Kingdom). The mass spectrometer was calibrated immediately before sample analysis using positive and negative ionisation calibration solution (recommended by Thermo Scientific). Additionally, the mass spectrometer scan rate was set at 4 Hz, giving a resolution of 25,000 (at 200 m/z) with a full-scan range of m/z 100 to 1,800 with continuous switching between positive and negative mode.

The PCA plot was obtained from lipid class percentage. The percentage of each lipid class was determined by combining all the lipid species within that class.

Elsewhere, lipid levels are expressed as a relative percentage abundance of each lipid on a per-line basis.

## QUANTIFICATION AND STATISTICAL ANALYSIS

Statistical analyses were performed in GraphPad Prism version 10 unless otherwise stated. Data are presented as mean ± SEM unless specified.

## List of Supplementary Materials

Figs. S1 to S4

Table S1

## Supplementary figure legends

**Figure S1. Characterisation of phenotypic screening in patient iPSC-DANs**. (A) Representative immunofluorescence images of iPSC-DANs cultured in 96- and 384-well formats and stained for TH/FOXA2 and MAP2/DAPI. (B) Percentage of TH-positive cells in Ctrl and *GBA-NS* cultures in 96- and 384-well formats; paired points indicate matched lines. (C) Coefficient of variation (CoV) for replicate compound wells and DMSO wells.

**Figure S2. Genetic and lipidomic validation of FASN and additional characterisation of FASN1 treatment.** (A) FASN mRNA expression in i3N transduced with non-targeting (NT) or FASN-targeting sgRNAs, expressed relative to NT controls. (B) Representative Western blot of FASN protein in i3N transduced with NT or FASN sgRNAs; β-actin was used as a loading control. (C) Quantification of FASN protein levels from Western blot analysis, expressed relative to NT controls. (D) sgRNA transduction efficiency in i3N, quantified as the percentage of mTagBFP2-positive cells in NT sgRNA-, FASN sgRNA-transduced and untransduced (UT) conditions. (E) PB-CAG-dCas9-KRAB-EF1α-Hygro-T2A-mNeonGreen construct used to generate the stable CRISPRi iPSC line. (F) Percentage of TH-positive and mNeonGreen-positive cells following differentiation of dCas9 iPSCs into dopaminergic neurons (iPSC-DANs). (H) Representative immunofluorescence image of differentiated dCas9 iPSC-DANs showing TH (orange) and mNeonGreen (green). Scale bar as shown. (I) Representative Western blot of FASN protein in dCas9 iPSC-DANs transduced with NT or FASN sgRNAs; β-actin was used as a loading control. (J) Quantification of FASN protein levels from Western blot analysis, expressed relative to NT controls. (K) sgRNA transduction efficiency in dCas9 iPSC-DANs, quantified as the percentage of mTagBFP2-positive cells in NT sgRNA-, FASN sgRNA-transduced and UT conditions. (L) Cell-toxicity signal following FASN1 treatment at the indicated concentrations, shown as fold change relative to DMSO; a positive control is shown for comparison. (M) Lipidomic analysis of Ctrl and *GBA-NS* iPSC-DANs following FASN1 treatment. Principal component analysis shows lipidomic profiles following treatment with DMSO or FASN1 (1, 5 or 10 µM). Pie charts show the relative abundance of lipid classes in Ctrl and *GBA-NS* iPSC-DANs treated with DMSO or FASN1 (5 µM). Data are shown as mean ± SEM where applicable. *P* < 0.05; ns, not significant.

**Figure S3. Validation of lysosomal GCase activity measurements in iPSC-DANs.** (A) Schematic representation of MDW941, a fluorescent activity-based probe that selectively labels active GCase. (B) MDW941 corrected spot intensity in Ctrl and *GBA-NS* neurons under baseline conditions and following treatment with the GCase inhibitor CBE as a negative control. (C) MDW941-positive spot number normalised to cell area in Ctrl and *GBA-NS* neurons under baseline conditions and following CBE treatment. (D) Expression of CLEAR genes in Ctrl and *GBA-NS* iPSC-DANs treated with DMSO or FASN1 (5 µM). Each point represents an individual line averaged across independent differentiations; data are mean ± SEM.

**Figure S4. FASN inhibition activates SREBP1 signalling and demonstrates *in vivo* target engagement**. (A) *SREBF1* mRNA expression in iPSC-DANs overexpressing SREBP1a or SREBP1c compared with control vector. (B) Representative Western blot of SREBP1 protein in iPSC-DANs overexpressing SREBP1a or SREBP1c compared with control vector; β-actin was used as a loading control. (C) Quantification of FASN protein levels from Western blot analysis, expressed relative to control vector. (D) BI 99179 plasma concentration over 24 h in control mice. (E) SNCA-OVX mice weight over time after dosing with BI 99179.

## Acknowledgements

We thank G. Onal, A. Lavayssiere, F. Yang, S. Gibson and N. Ibarra-Aizpurua for expert assistance in stem cell culture and neuronal differentiation. We thank C. Lang and A. Mehta for providing human brain-derived cDNA. We thank K. Blachowiak for expert assistance in mouse husbandry, drug dosing and tissue collection.

## Author contributions

M.C.C., J.B., R.M.G., E.B., U.C., W.M., S.H., P.S. J.H., M.T. and B.R. designed, performed, and analysed experimental research. B.J. performed lipidomics analysis, supervised by A.K.. D.B., A.D. and D.S. provided access and advice on the use of the small molecule library and analogues. R.W.-M. and B.R. devised and supervised the project. M.C.C, B.R. and R.W.-M. wrote the paper. All authors reviewed the manuscript and approved its submission.

## Data and materials availability

All data, code and materials used in the analysis will be made available to researchers, subject to the usual materials transfer agreement (MTA) requirements, through contacting the lead author. Lipdomics datasets have been uploaded and are available from Zenodo (doi: 10.5281/zenodo.20185957). Permutation analysis was performed using a custom Python notebook, publicly available at https://github.com/mariaclaudiacaia/plate-reader-assays/blob/ac931a409d816715538ed043849087293e134f78/permutation_test.ipynb.

