## Supplemental Tables for "Phenotypic screening in Parkinson’s iPSC-dopamine neurons identifies the FASN-SREBP1 axis as a potential therapeutic target"

| Compound ID | Primary target |
|---|---|
| 3555 | H3 antagonist (inactive) |
| 7175 | IKACh (potential artefact) |
| 8466 | ATM |
| 1733 | MTH1 |
| 6225 | Tyrosine kinases |
| 7732 | D1 activator/potentiator |
| 0497 | PKC |
| 7386 | MEK; ERK |
| 6122 | FASN |
| 2930 | mGLuR2 |
| 9844 | PKC |
| 8891 | PKC |
| 6096 | Protease inhibitor |
| 5581 | PAK1 |
| 9259 | D2/NET (inactive) |
| 9328 | MEK1 |
| 9128 | BACE |
| 7725 | PIP5K1C |
| 0513 | ERK1 |
| 0300 | EGFR |
| 1446 | ATM |
| 8558 | FASN |
| 9365 | Pan CDK |

**Supplemental Table 1**
**Hits from primary screen.**

| Primer | Target | Sequence |
|---|---|---|
| **ACACA-F** | ACACA | TGATTTGGGGATCTCTAGCC |
| **ACACA-R** | ACACA | CTGGAGAAGCCACAGTGAAA |
| **ACLY-F** | ACLY | TCTTTGTCCGAAGAGGTGGC |
| **ACLY-R** | ACLY | GCCAAAGACATGGATGGGGA |
| **ACSS2-F** | ACSS2 | GTGGAGTCAGCACTTGTGGA |
| **ACSS2-R** | ACSS2 | GGCATTCACCCTTCACAGGA |
| **ACTIN-F** | ATCB | GGGGTGTTGAAGGTCTCAAA |
| **ACTIN-R** | ATCB | AGAAAATCTGGCACCACACC |
| **CTSD-F** | CTSD | GCAAATGCTGCGACATCGCTTG |
| **CTSD-R** | CTSD | GCCATAGTGGATGTGAAAACGAGG |
| **CYC1-F** | CYC1 | CAGGGAAGCTGTTCGACTATTT |
| **CYC1-R** | CYC1 | TGTGCCGCTTTATGGTGTAG |
| **DHCR24-F** | DHCR24 | CAGGAGAACCACTTCGTGGAAG |
| **DHCR24-R** | DHCR24 | CCACATGCTTAAAGAACCACGGC |
| **ERO1A-F** | ERO1A | AATGTCGTCTGTGGGGAAAG |
| **ERO1A-R** | ERO1A | AGGTCCACTTTCTGGCATATTT |
| **FASN-F** | FASN | AGATGGCTTGCTGGAGAACC |
| **FASN-R** | FASN | AAGCCGTAGTTGCTCTGTCC |
| **GAPDH-F** | GAPDH | CTGACTTCAACAGCGACACC |
| **GAPDH-R** | GAPDH | ATGAGGTCCACCACCCTG |
| **GBA-F** | GBA | CTGCTGCTCTCAACATCTCTTGCC |
| **GBA-R** | GBA | CTAGGTGCGGATGGAAGAAGTCAC |
| **HMGCR-F** | HMGCR | GACGTGAACCTATGCTGGTCAG |
| **HMGCR-R** | HMGCR | GGTATCTGTTTCAGCCACTAAGG |
| **HPRT1-F** | HPRT1 | GCCAGACTTTGTTGGATTTG |
| **HPRT1-R** | HPRT1 | CTCTCATCTTAGGCTTTGTATTTTG |
| **ITPR1-F** | ITPR1 | GTGACAGGAAACATGCAGACTCG |
| **ITPR1-R** | ITPR1 | CAGCAGTTGCACAAAGACAGGC |
| **LAMP1-F** | LAMP1 | CGTGTCAAGGAAGGCTTTTCAG |
| **LAMP1-R** | LAMP1 | CTGTTCTCGTCCAGCAGACACT |
| **LDLR-F** | LDLR | CCCAGCGAAGATGCGAAGATAT |
| **LDLR-R** | LDLR | GGTGAAGAAGAGGTAGGCGATG |
| **LPIN1-F** | LPIN1 | CCACTCTTGCCCATGATCGA |
| **LPIN1-R** | LPIN1 | ACCAAGATGTCGGCTTCGTT |
| **MCOLN1-F** | MCOLN1 | GACAAGTTTCGAGCCAAAGGG |
| **MCOLN1-R** | MCOLN1 | GATGGTGTTTCTTCTTCCCGGA |
| **RPL13-F** | RPL13 | CCTGGAGGAGAAGAGGAAAGAGA |
| **RPL13-R** | RPL13 | TTGAGGACCTCTGTGTATTTGTCAA |
| **SCARB2-F** | SCARB2 | TTGATAAAGAGACGGCGAGTC |
| **SCARB2-R** | SCARB2 | ACCAAGCAAAGGCAATGTTT |
| **SQLE-F** | SQLE | CCTGAATCAGAAAATAAGGAGCA |
| **SQLE-R** | SQLE | GCTTGTTTCTGAAATATTGGTTCC |
| **SREBF1-F** | SREBF1 | TCCACCCCTGTGTTAGGCTA |
| **SREBF1-R** | SREBF1 | ACACTCTGGACCTGGGTGT |
| **SREBF2-F** | SREBF2 | TGGGAGAGTTCCCTGACTTG |
| **SREBF2-R** | SREBF2 | CACTGAAGGGTCCACAGCTC |

**Supplemental Table 2**
**Primer list.**
