## Supplemental Figures for "Phenotypic screening in Parkinson’s iPSC-dopamine neurons identifies the FASN-SREBP1 axis as a potential therapeutic target"

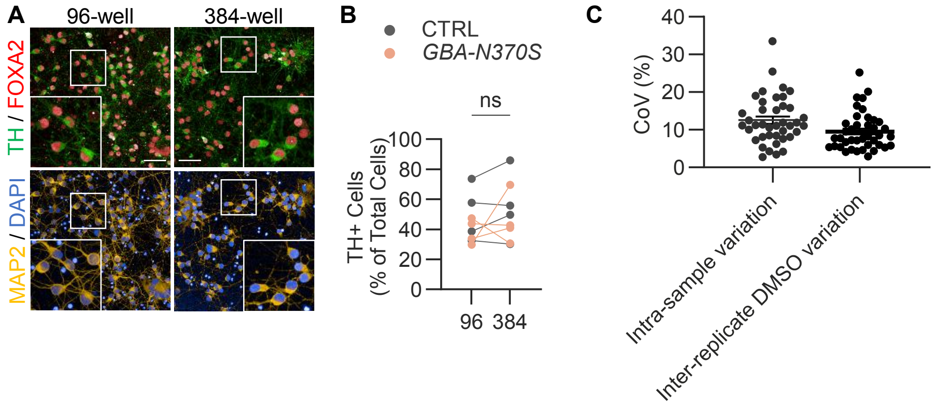
A
96-well
384-well
TH / FOXA2
MAP2 / DAPI
B
CTRL
GBA-N370S
ns
TH+ Cells
(% of Total Cells)
100
80
60
40
20
0
96
384
C
CoV (%)
40
30
20
10
0
Intra-sample variation
Inter-replicate DMSO variation

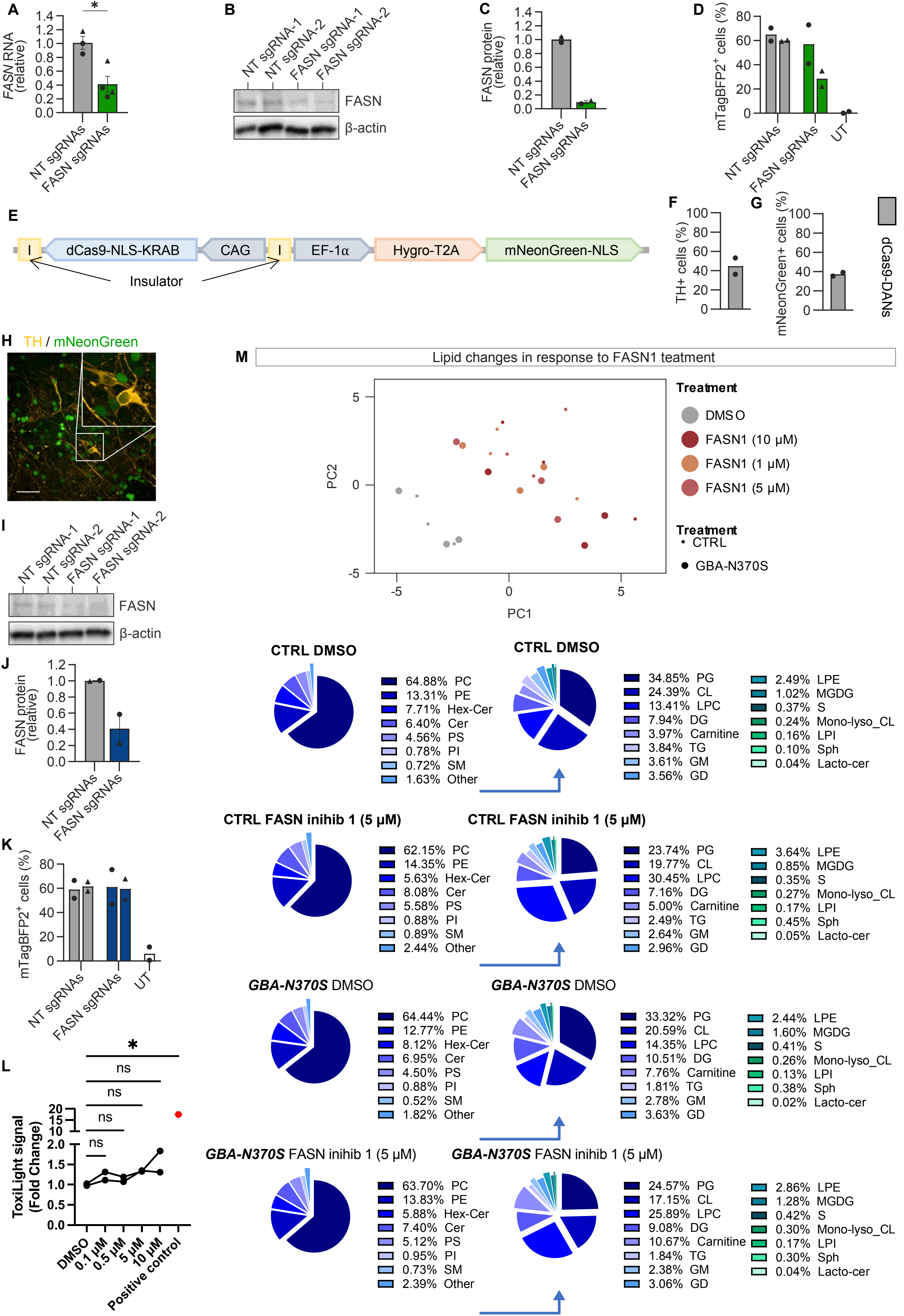

A
FASN RNA (relative)
*
NT sgRNAs
FASN sgRNAs
B
NT sgRNA-1
NT sgRNA-2
FASN sgRNA-1
FASN sgRNA-2
FASN
β-actin
C
FASN protein (relative)
NT sgRNAs
FASN sgRNAs
D
mTagBFP2+ cells (%)
NT sgRNAs
FASN sgRNAs
UT
E
I
dCas9-NLS-KRAB
CAG
I
EF-1α
Hygro-T2A
mNeonGreen-NLS
Insulator
F
TH+ cells (%)
G
mNeonGreen+ cells (%)
dCas9-DANs
H
TH / mNeonGreen
I
NT sgRNA-1
NT sgRNA-2
FASN sgRNA-1
FASN sgRNA-2
FASN
β-actin
J
FASN protein (relative)
NT sgRNAs
FASN sgRNAs
K
mTagBFP2+ cells (%)
NT sgRNAs
FASN sgRNAs
UT
L
ToxiLight signal (Fold Change)
*
ns
ns
ns
ns
DMSO
0.1 µM
0.5 µM
5 µM
10 µM
Positive control
M
Lipid changes in response to FASN1 teatment
PC2
PC1
Treatment
DMS O
FASN1 (10 µM)
FASN1 (1 µM)
FASN1 (5 µM)
Treatment
CTRL
GBA-N370S
CTRL DMSO
64.88% PC
13.31% PE
7.71% Hex-Cer
6.40% Cer
4.56% PS
0.78% PI
0.72% SM
1.63% Other
CTRL DMSO
34.85% PG
24.39% CL
13.41% LPC
7.94% DG
3.97% Carnitine
3.84% TG
3.61% GM
3.56% GD
2.49% LPE
1.02% MGDG
0.37% S
0.24% Mono-lyso_CL
0.16% LPI
0.10% Sph
0.04% Lacto-cer
CTRL FASN inihib 1 (5 µM)
62.15% PC
14.35% PE
5.63% Hex-Cer
8.08% Cer
5.58% PS
0.88% PI
0.89% SM
2.44% Other
CTRL FASN inihib 1 (5 µM)
23.74% PG
19.77% CL
30.45% LPC
7.16% DG
5.00% Carnitine
2.49% TG
2.64% GM
2.96% GD
3.64% LPE
0.85% MGDG
0.35% S
0.27% Mono-lyso_CL
0.17% LPI
0.45% Sph
0.05% Lacto-cer
GBA-N370S DMSO
64.44% PC
12.77% PE
8.12% Hex-Cer
6.95% Cer
4.50% PS
0.88% PI
0.52% SM
1.82% Other
GBA-N370S DMSO
33.32% PG
20.59% CL
14.35% LPC
10.51% DG
7.76% Carnitine
1.81% TG
2.78% GM
3.63% GD
2.44% LPE
1.60% MGDG
0.41% S
0.26% Mono-lyso_CL
0.13% LPI
0.38% Sph
0.02% Lacto-cer
GBA-N370S FASN inihib 1 (5 µM)
63.70% PC
13.83% PE
5.88% Hex-Cer
7.40% Cer
5.12% PS
0.95% PI
0.73% SM
2.39% Other
GBA-N370S FASN inihib 1 (5 µM)
24.57% PG
17.15% CL
25.89% LPC
9.08% DG
10.67% Carnitine
1.84% TG
2.38% GM
3.06% GD
2.86% LPE
1.28% MGDG
0.42% S
0.30% Mono-lyso_CL
0.17% LPI
0.30% Sph
0.04% Lacto-cer

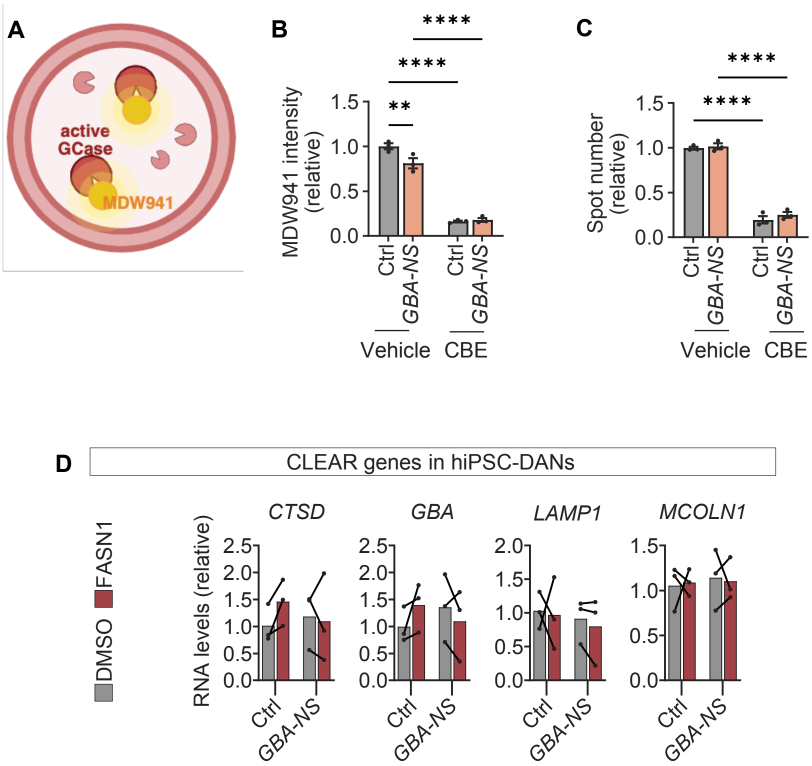
A
active
GCase
MDW941
B
MDW941 intensity
(relative)
1.5
1.0
0.5
0.0
Ctrl
GBA-NS
Ctrl
GBA-NS
Vehicle
CBE
**
****
****
C
Spot number
(relative)
1.5
1.0
0.5
0.0
Ctrl
GBA-NS
Ctrl
GBA-NS
Vehicle
CBE
****
****
D
CLEAR genes in hiPSC-DANs
DMSO
FASN1
RNA levels (relative)
CTSD
GBA
LAMP1
MCOLN1
2.5
2.0
1.5
1.0
0.5
0.0
Ctrl
GBA-NS

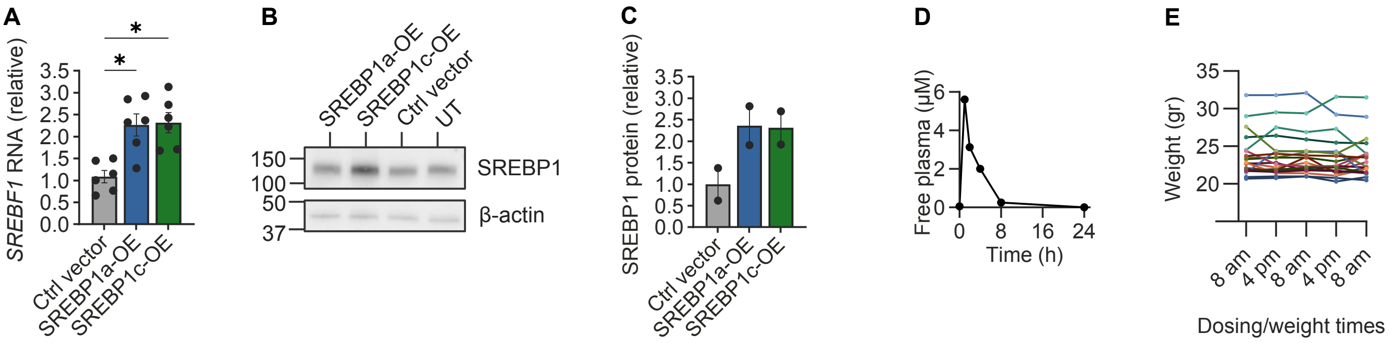

A
*
*
SREBF1 RNA (relative)
3.5
3.0
2.5
2.0
1.5
1.0
0.5
0.0
Ctrl vector
SREBP1a-OE
SREBP1c-OE
B
SREBP1a-OE
SREBP1c-OE
Ctrl vector
UT
150
100
50
37
SREBP1
β-actin
C
SREBP1 protein (relative)
3.5
3.0
2.5
2.0
1.5
1.0
0.5
0.0
Ctrl vector
SREBP1a-OE
SREBP1c-OE
D
Free plasma (μM)
6
4
2
0
0
8
16
24
Time (h)
E
Weight (gr)
35
30
25
20
8 am
4 pm
8 am
4 pm
8 am
Dosing/weight times
